# Comprehensive prediction of robust synthetic lethality between paralog pairs in cancer cell lines

**DOI:** 10.1101/2020.12.16.423022

**Authors:** Barbara De Kegel, Niall Quinn, Nicola A. Thompson, David J. Adams, Colm J. Ryan

## Abstract

Pairs of paralogs may share common functionality and hence display synthetic lethal interactions. As the majority of human genes have an identifiable paralog, exploiting synthetic lethality between paralogs may be a broadly applicable approach for targeting gene loss in cancer. However only a biased subset of human paralog pairs has been tested for synthetic lethality to date. Here, by analysing genome-wide CRISPR screens and molecular profiles of over 700 cancer cell lines, we identify features predictive of synthetic lethality between paralogs, including shared protein-protein interactions and evolutionary conservation. We develop a machine-learning classifier based on these features to predict which paralog pairs are most likely to be synthetic lethal and to explain why. We show that our classifier accurately predicts the results of combinatorial CRISPR screens in cancer cell lines and furthermore can distinguish pairs that are synthetic lethal in multiple cell lines from those that are cell-line specific.

## Introduction

Gene duplication is the primary means by which new genes are created (Zhang, 2003) and consequently the majority of human genes are duplicates (paralogs) (Zerbino et al., 2018). Although paralog pairs may functionally diverge over time, many maintain at least some functional overlap, even after long evolutionary periods (Conant and Wolfe, 2008; Vavouri et al., 2009). This functional overlap may allow paralog pairs to buffer each other’s loss, contributing to the overall ability of cells and organisms to tolerate genetic perturbations. Indirect evidence for the contribution of paralog buffering to genetic robustness is provided by the finding that, in both model organisms and cancer cell lines, paralog genes are less likely to be essential than genes with no identifiable paralogs (Blomen et al., 2015; Dandage and Landry, 2019; De Kegel and Ryan, 2019; Gu et al., 2003; Wang et al., 2015). More direct evidence for functional buffering is provided by observations of synthetic lethality between paralog pairs (Dean et al., 2008; Dede et al., 2020; De Kegel and Ryan, 2019; DeLuna et al., 2008; VanderSluis et al., 2010). Synthetic lethality occurs when the perturbation of two genes individually is well tolerated but their combined perturbation results in a loss of cellular fitness or cell death. Synthetic lethal relationships appear to be much more frequent among paralog pairs than other gene pairs – in the budding yeast *Saccharomyces cerevisiae*, where comprehensive double perturbation screens have been performed, 25-35% of paralog pairs are synthetic lethal, compared to <5% of random gene pairs (Costanzo et al., 2016; Dean et al., 2008; DeLuna et al., 2008; Musso et al., 2008). It is not yet established how frequent synthetic lethality is among paralog pairs in human cells.

As gene loss is a common feature of tumour cells, synthetic lethality has long been proposed as a means of selectively killing tumour cells while limiting toxicity to healthy, non-transformed cells (Hartwell et al., 1997; Kaelin, 2005; O’Neil et al., 2017). The first therapies that exploit synthetic lethality – the use of PARP inhibitors in patients with loss-of-function *BRCA1* or *BRCA2* mutations – have now been approved for clinical use in select breast, ovarian, and prostate tumours (Lord and Ashworth, 2017). Many large-scale efforts are now focussed on the identification of new synthetic lethal treatments, in particular for cancers with recurrent genetic alterations that currently have no effective targeted therapies associated with them (Behan et al., 2019; Han et al., 2017; Huang et al., 2020). However, it remains challenging to identify new, robust synthetic lethal interactions in cancer due to a lack of understanding of what makes a gene pair likely to be synthetic lethal.

As >60% of human genes have an identifiable paralog, and as paralog pairs are more likely to be synthetic lethal than random gene pairs, exploiting synthetic lethality among paralog pairs may be a widely applicable approach for targeting a variety of gene loss events in cancer. This includes targeting recurrently altered driver genes, such as the synthetic lethal interaction between the tumour suppressor *SMARCA4* and its paralog *SMARCA2*, and targeting passenger gene loss events, such as the synthetic lethal interaction between *ME2* and its paralog *ME3*. *ME2* is not believed to be a cancer driver gene but is located in the same chromosomal region as the tumour suppressor *SMAD4* and is frequently homozygously deleted in tumours that have lost *SMAD4* (Dey et al., 2017). Overall the identification of synthetic lethal interactions involving paralogs that are recurrently lost in cancer has resulted in about a dozen validated paralog synthetic lethalities (Benedetti et al., 2017; Dey et al., 2017; Ehrenhöfer-Wölfer et al., 2019; Helming et al., 2014; Hoffman et al., 2014; van der Lelij et al., 2020; Lord et al., 2020; Muller et al., 2012; Ogiwara et al., 2016; Oike et al., 2013; Schick et al., 2019; Szymańska et al., 2020; Tsherniak et al., 2017; Viswanathan et al., 2018).

To date, synthetic lethal interactions between paralog pairs have primarily been discovered through the analysis of large scale loss-of-function screens (e.g. *ARID1A* mutant cell lines were found to be sensitive to loss of its paralog *ARID1B* through analysis of genome-wide shRNA screens of 165 tumour cell lines (Helming et al., 2014)) or through the rational testing of individual paralog pairs (*ME3* was tested for synthetic lethality with *ME2* because of an understanding of their role in metabolism (Dey et al., 2017)). Recently combinatorial screening using CRISPR gene perturbation has emerged as an alternative approach for determining synthetic lethality between paralog pairs (Dede et al., 2020; Gonatopoulos-Pournatzis et al., 2020; Parrish et al., 2020; Thompson et al., 2021). In brief, these screens use individual and paired guide RNAs to determine the expected and observed fitness consequences of perturbing each gene pair. Synthetic lethality can then be identified where the observed fitness defect is significantly greater than that expected from the fitness defects induced by the individual guide RNAs. This approach allows researchers to quantify synthetic lethal relationships between hundreds of gene pairs in pooled screens. However, there are tens of thousands of paralog pairs in the human genome (Zerbino et al., 2018) and to date only ∼5% of them have been tested for synthetic lethality in combinatorial screens. Furthermore, many of the reported hits from these screens have been identified in a cell-line specific fashion (Dede et al., 2020; Gonatopoulos-Pournatzis et al., 2020; Parrish et al., 2020; Thompson et al., 2021) and therefore may not be useful for the development of targeted therapeutics where genetic heterogeneity within and between tumours is a major challenge (Henkel et al., 2019; Lord et al., 2020; Ryan et al., 2018). There is thus a need for a computational approach to identify which of the tens of thousands of paralog pairs are most likely to be ‘robust’ synthetic lethals that operate across multiple genetic backgrounds.

A number of statistical and machine learning approaches have been developed for genome-wide prediction of synthetic lethality (Benstead-Hume et al., 2019; Cai et al., 2020; Jacunski et al., 2015; Lee et al., 2018) but these have suffered from the lack of a set of validated synthetic lethal interactions for training and evaluation. Furthermore, due their specific input requirements, most of these approaches can only generate predictions for a limited number of paralog pairs. There are as of yet no approaches that are tailored specifically to predict synthetic lethality between paralogs. We, and others, have previously identified a small number of features (amino acid sequence identity, duplication mode and protein complex membership) that are somewhat predictive of synthetic lethality between paralog pairs (De Kegel and Ryan, 2019; DeLuna et al., 2008; Guan et al., 2007) but a more comprehensive analysis of what makes a paralog pair likely to be synthetic lethal has yet to be performed.

Here we develop an ensemble classifier to predict synthetic lethality among paralog pairs in cancer cell lines and to provide explanations for why specific paralog pairs are likely to be synthetic lethal (SL). We first show that existing combinatorial screens have severe selection bias, in terms of the pairs that are included for screening, and that the hits from these screens are primarily cell line specific. To address these limitations, we derive a dataset of SL (and non-SL) paralog pairs by integrating molecular profiling data with genome-wide CRISPR screens in 762 heterogeneous cancer cell lines from the DepMap project (Meyers et al., 2017). Because these SLs are identified across multiple cell lines they are more likely to be robust to genetic heterogeneity and unlikely to be cell line specific effects. We then assess the ability of 22 different features of paralog pairs to distinguish SL from non-SL pairs in this dataset. We identify several individual features, such as shared protein-protein interactions and evolutionary conservation, that are predictive of synthetic lethal relationships. We integrate all features using a random forest classifier and show that, for our dataset of paralog pairs, this classifier can greatly outperform predictions based on the best individual feature. We further demonstrate that our classifier can accurately predict the consensus results from four independent combinatorial CRISPR screens. In addition, we show that our predictions can distinguish between cell-line specific and robust SLs. Finally, we make predictions for all paralog pairs, accompanied by local explanations that indicate why specific pairs are likely to be SL.

## Results

### Combinatorial screens have severe bias and primarily identify cell line specific SLs

To understand what makes a paralog pair more, or less, likely to be synthetic lethal, one needs a training dataset of ‘true positive’ (SL) and ‘true negative’ (non-SL) pairs. The number of SL interactions between paralogs with low-throughput experimental validation in human cells is extremely small – from the literature we identified only 12 such paralog pairs (Table S1). This small set of ‘true positives’ precludes any meaningful analysis of what makes a paralog pair likely to be SL. Combinatorial CRISPR screens of paralog pairs, which have assayed hundreds of paralog pairs in pooled screens, may provide a more comprehensive set of true positives and negatives. However, these screens have selected paralogs for inclusion in a biased fashion and we anticipated that this bias might also limit their use as training data.

We obtained data from four independent combinatorial screening efforts – two used a CRISPR/Cas9 approach (Parrish et al., 2020; Thompson et al., 2021), one used a CRISPR/enCas12a approach (Dede et al., 2020), and one used a hybrid Cas9-Cas12a approach called CHyMErA (Gonatopoulos-Pournatzis et al., 2020) (see Methods). These four screens each tested several hundred (∼400-1000) paralog pairs which were selected for inclusion according to different criteria (e.g. pairs with high co-expression, pairs from whole genome duplication events). To understand the consequences of these inclusion criteria we compared the properties of paralogs included in these screens to an unbiased reference set of ∼36.6k protein-coding paralog pairs obtained from Ensembl (Zerbino et al., 2018) (see Methods). We find that in all screens there is severe bias in the selected paralog pairs and that they are not representative of paralog pairs in general (Fig. 1A-D). In comparison to the reference set, the paralog pairs selected for each screen share higher sequence identity (Fig. 1A), tend to come from smaller families (Fig. 1B), are more likely to be whole genome duplicates (Fig. 1C) and are more likely to be each other’s closest (i.e. most sequence similar) paralog (Fig. 1D). The differences are stark: the median family size for all screened pairs is only 3, while for all paralog pairs it is 11; ∼73% of screened pairs are labelled as whole genome duplicates (WGDs), compared to only ∼19% of all paralog pairs; and ∼89% of screened pairs are closest pairs, compared to only ∼13% of all paralog pairs. While this extreme bias is not surprising, given that the screens attempted to select pairs likely to be SL, it impedes analysis of what features make a paralog pair more or less likely to be SL. For example, we cannot learn what predicts SL among non-closest paralog pairs if the available data consists almost entirely of closest pairs.

**Figure 1.**
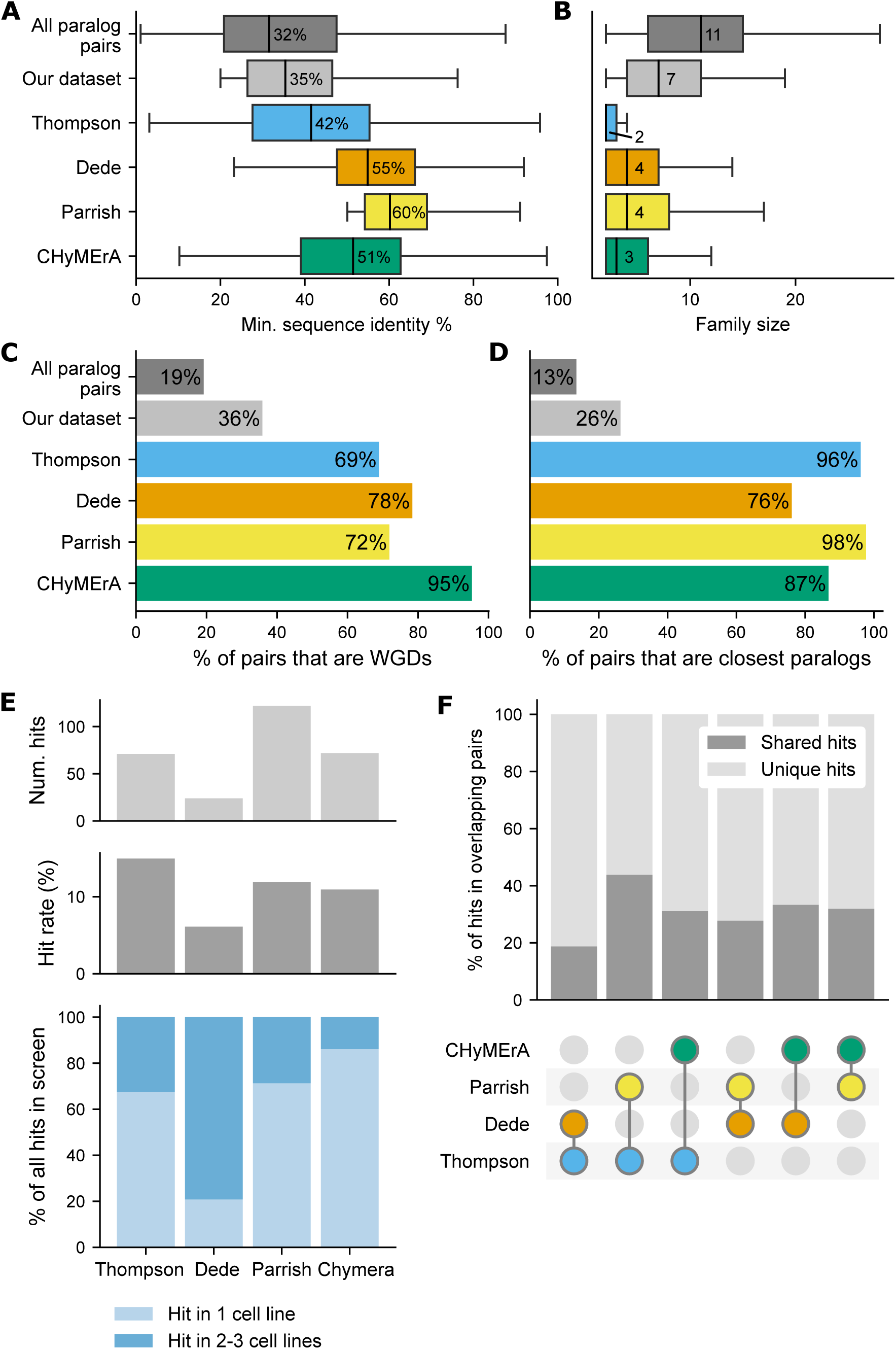
Selection bias and cell-line specific hits in four combinatorial screens of paralog pairs. (A,B) Boxplots showing the distribution of (A) minimum amino acid sequence identity and (B) family size for: all paralog pairs (dark grey), paralog pairs in our training dataset (light grey, see Fig. 2A), and paralog pairs in each of four combinatorial screens (colored). For each box plot, the central line represents the median (also shown as annotation), the top and bottom represent the 1st and 3rd quartile, and the whiskers extend to 1.5 times the interquartile range past the box. Outliers are not shown. (C,D) Bar charts showing the percentage of paralogs pairs that are (C) whole genome duplicates (WGDs) and (D) closest paralogs for the same six datasets of paralog pairs. (E) Top: bar charts showing the number and percentage of tested paralog pairs that were identified as SL in any cell line in each screen. Bottom: bar chart showing the percentage of hits that were observed to be cell-line specific vs. reproduced in multiple cell lines. (F) For each pair of screens (as indicated) the percentage of hits among pairs tested in both screens that are shared (i.e. hits in both screens) vs. unique (i.e. hits in only one screen).

Each of the four studies assayed two or three different cell lines from different lineages (see Methods). Where available, we obtained the authors reported SL hits for each study. The CHyMErA study (Gonatopoulos-Pournatzis et al., 2020) reported a broader set of negative genetic interaction hits which we filtered to only include synthetic lethals (see Methods). We observe that all four screens have a fairly consistent hit rate of ∼6-15% (median ∼11%, Fig. 1E, top) but that many of the hits in each screen are cell line specific (Fig. 1E, bottom). The majority (∼68-86%) of SL pairs identified in the Thompson, Parrish and CHyMErA screens are hits in only a single cell line. Exceptionally, a large percentage (∼79%) of the hits in the Dede *et al*. screen are reproduced in at least two of the three cell lines assayed. This screen had the lowest hit rate (∼6%), so more stringent hit-calling might partially explain why there is higher concordance between cell lines.

Through pairwise comparisons of the four screens we find that the overlap of hits between screens is moderate (Fig. 1F). For each pair of screens we considered only the paralog pairs that were tested in both screens and which were a hit in either screen. Using this approach we determined that on average ∼31% of the hits are shared, while the rest are unique to one of the two screens. The Thompson and Dede screens have the lowest percentage of shared hits (∼19%) while the Thompson and Parrish screens have the highest (∼44%). The overlap of (non-)hits among paralog pairs tested in both screens is significant in all cases (Fisher’s exact test, *p*<=0.0001), except for the Thompson-Dede comparison (Fisher’s exact test, *p*=0.06).

Only a single cell line, RPE-1, was screened in more than one study (Thompson *et al*. and Gonatopoulos-Pournatzis *et al*. (CHyMErA)). For the hits in this cell line we observe very poor agreement between the two screens (Fig S1C); among the 26 paralog pairs that were screened by both and hits in either screen, only 2 (∼8%) are hits in both screens. This suggests that the observed screen specificity of hits is likely due to a combination of cell line specific hits and false positive and false negative hits.

### Identification of a set of robust synthetic lethal paralog pairs from genome-wide screens

Consistent with our analysis of hits from combinatorial screens, it has become increasingly clear that many synthetic lethal interactions operate only within specific genetic backgrounds or contexts (Henkel et al., 2019; Ku et al., 2020; Lord et al., 2020; Ryan et al., 2018). This has significant consequences for the development of therapeutics based on SL interactions – interactions that are observed in multiple cell lines are more likely to be robust in the face of the molecular heterogeneity that exists within and between tumours and consequently may make more promising drug targets (Ryan et al., 2018). As many of the hits from the existing combinatorial screens of paralog pairs appear to be context-dependent, and, as shown, there is severe bias in the paralog pairs selected for screening, we cannot reliably use the results of these screens to investigate features of robust SL paralog pairs. We therefore used a computational approach to identify an unbiased set of robust SL and non- SL paralog pairs from existing whole genome CRISPR screening data (Fig. 2A).

**Figure 2.**
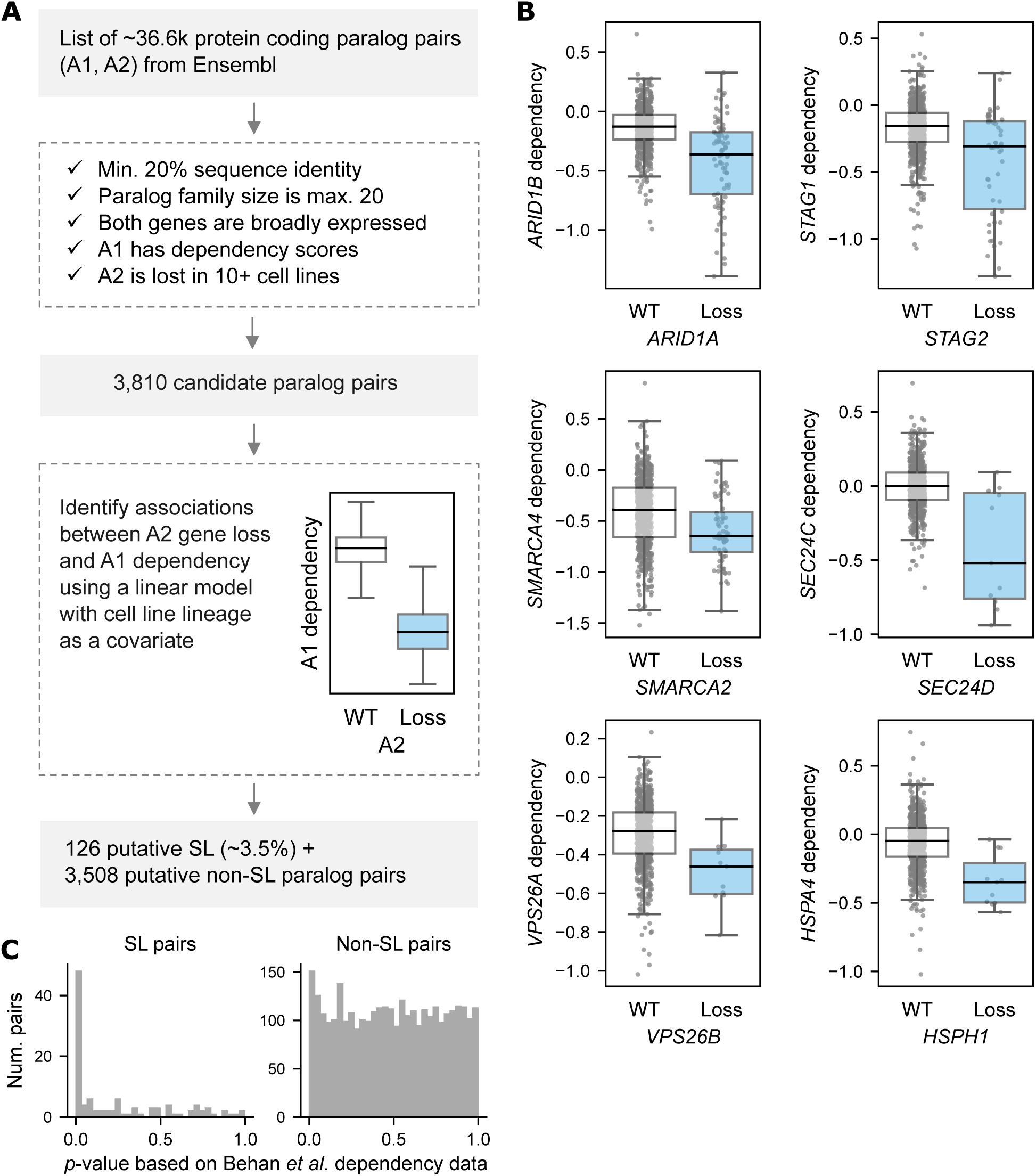
Identifying a dataset of robust SL paralog pairs. (A) Workflow of our computational approach to identify a set of robust SL paralog pairs from the reference list of Ensembl protein-coding paralog pairs. The two genes that make up each paralog pair are referred to as A1 and A2. (B) Boxplots showing A1 dependency (CERES score: larger negative score implies larger negative LFC and thus increased dependency) in cell lines where A2 is lost (blue box) vs. where A2 is wild-type (white box) for six of the SL paralog pairs that we identified using the approach in (A). For each box plot, the black central line represents the median, the top and bottom represent the 1st and 3rd quartile, and the whiskers extend 1.5 times the interquartile range past the box. Each dot represents one cell line in which A1 was screened. (C) Histograms showing, for our SL and non-SL paralog pairs, the distribution of *p*-values obtained from testing for association between A1 dependency and A2 gene loss using dependency data from Behan *et al*.

Genome-wide CRISPR screens report on the fitness cost of perturbing each protein-coding gene in a given cell line and, when combined with molecular profiling data, can provide insight into which genes are essential in cell lines with specific alterations. We integrated genome-wide CRISPR screens of 762 heterogeneous cancer cell lines (Meyers et al., 2017) (DepMap 20Q2, Broad) with molecular profiling data for the same cell lines (Ghandi et al., 2019) and searched for cases where loss or silencing of one member of a paralog pair (termed A2) results in increased dependency on the other member (termed A1) (Fig. 2A). As in previous work (De Kegel and Ryan, 2019) we first reprocessed the DepMap CRISPR screen results to remove guides that might target multiple genes, an issue that disproportionately affects sequence-similar paralogs and which could confound our results (Fortin et al., 2019) (Table S2, see Methods). We started our analysis with the reference list of paralog pairs obtained from Ensembl (Zerbino et al., 2018) and filtered this list according to a number of criteria (Fig. 2A). Briefly, we removed pairs with very low (<20%) amino acid sequence identity, pairs from very large paralog families (>20 genes), pairs with tissue- specific expression, and pairs that were not present in our reprocessed DepMap CRISPR screen data (see Methods). Finally, we required that at least one gene in the pair is lost – i.e. subject to a homozygous deletion, loss-of-function mutation, or loss of mRNA expression – in at least 10 (and max. 90%) of the 762 cancer cell lines (Table S3, see Methods). The application of all five filtering steps culminated in a set of 3,810 candidate paralog pairs for further analysis (Fig. 2A).

To identify associations between loss of one paralog (A2) and altered dependency upon the other paralog (A1) among these candidate pairs we used a multiple linear regression model with A1 dependency as the dependent variable, and A2 status (wildtype or loss) and cell line lineage as the independent variables (see Methods). Cell line lineage was included to account for variation in A1 dependency that is due to tissue type. For 126 of the 3,810 candidate pairs we found that A2 status was significantly associated with A1 dependency (10% FDR), i.e. A2 loss resulted in increased A1 dependency (lower A1 fitness score), indicative of a synthetic lethal relationship (Table S4). These putative SL pairs include six of the previously validated synthetic lethalities that we curated from the literature (Table S1), including *ARID1A/ARID1B (Helming et al., 2014), STAG1/STAG2* (van der Lelij et al., 2020) and *SMARCA2/SMARCA4* (Oike et al., 2013), as well as several pairs with strong associations that have not, to our knowledge, been previously reported, such as *SEC24C/SEC24D*, *VPS26A/VPS26B* and *HSPA4/HSPH1* (Fig. 2B).

For our robust (non-)SL dataset the 126 putative SL pairs were labelled as true positives, while the 3,508 pairs for which there was no association between A2 loss and A1 dependency (uncorrected *p*>0.05) were labelled as true negatives (Table S5). The remaining tested pairs (with uncorrected *p*≤0.05 but FDR>10%) were discarded. The final dataset thus contains 3,634 paralog pairs, of which ∼3.5% are considered SL. To validate this dataset we re-tested both the SL and non-SL pairs using the same linear regression model but with dependency scores for 242 cell lines from a set of independently performed whole genome CRISPR screens (Behan et al., 2019) (see Methods). While almost half (47.3%) of the SL pairs from our dataset had a significant association between A1 dependency and A2 loss when using the Behan *et al*. dependency data, none of the non-SL pairs did (Fig. 2C).

Comparing our dataset of SL and non-SL paralog pairs against the reference set of all paralog pairs we find that it is considerably less biased than the sets of paralog pairs tested in each of the existing combinatorial screens (Fig. 1A-D) (Dede et al., 2020; Gonatopoulos-Pournatzis et al., 2020; Parrish et al., 2020; Thompson et al., 2021). Given that our computational pipeline filtered on sequence identity and family size, our dataset still has somewhat different feature distributions than the reference set, but overall it is much more representative of all paralog pairs. We observe that there is moderate overlap between the hits in our dataset and the hits in each of the screens (Fig. S1D). Similar to what we saw for the screen-to-screen comparisons, on average ∼32% of the hits are shared. Thus our computational approach has identified several of the same hits as the combinatorial screens but against a much less biased background. In addition, as the SL pairs in our dataset were identified from combined analysis of heterogeneous cancer cell lines from 26 different lineages, they are expected to be largely context-agnostic, i.e. robust to cancer type and genetic background.

### Identification of features predictive of synthetic lethality between paralogs

We next sought to identify features that are predictive of synthetic lethality between paralogs. To this end we computed 22 features of paralog pairs (Table S6), including properties of the individual genes in the paralog pair (e.g. mean gene expression of the more highly expressed paralog), and comparative properties that quantify some relationship between the two genes (e.g. mRNA expression correlation). The features we assembled are either Boolean (True/False) or quantitative properties and fall into three broad categories: sequence, neighborhood and expression features (Table S6). Features in the ‘sequence’ category are derived from analyses of the protein-coding sequences of the pairs and include sequence identity, measures of conservation in other species, gene age, and the ratio of the lengths of the two proteins. ‘Neighborhood’ features pertain mainly to the protein-protein interactions (PPI) and the protein complex membership of the genes in each paralog pair. The ‘expression’ features are calculated from measurements of mRNA expression in non-diseased tissues (GTEx Consortium, 2020) and comprise the average expression and expression correlation of each paralog pair. Three of the features – sequence identity, protein complex membership and duplication mode (whole genome vs. small scale duplication) – are ones for which we previously observed an association with putative synthetic lethality in human cells (De Kegel and Ryan, 2019). In budding yeast, sequence identity and duplication mode have also been associated with paralog SL (DeLuna et al., 2008; Guan et al., 2007), while protein complex membership, protein complex essentiality, protein-protein interactions, co-localisation and co-expression have all been associated with negative genetic interactions in general (Bandyopadhyay et al., 2008; Costanzo et al., 2010, 2016; Michaut et al., 2011; Ryan et al., 2013).

We assessed each feature’s ability to predict synthetic lethality in our dataset of SL and non-SL paralog pairs by treating each feature as an individual classifier and computing area under the receiver operating characteristic curve (ROC AUC) and average precision (Fig. 3A, Table S6, see Methods). Average precision is a measure of the area under the precision-recall (PR) curve and is often a more sensitive measure of performance when a classification challenge has high class imbalance, i.e. when the positive examples are much rarer than the negative examples (Lever et al., 2016). As our dataset has high class imbalance (many more paralog pairs are not SL than are SL) we considered both metrics for evaluation. It should be noted that by chance the average precision is only ∼3.5%, and thus average precision is expected to be small (in absolute terms) even for well-performing classifiers.

**Figure 3.**
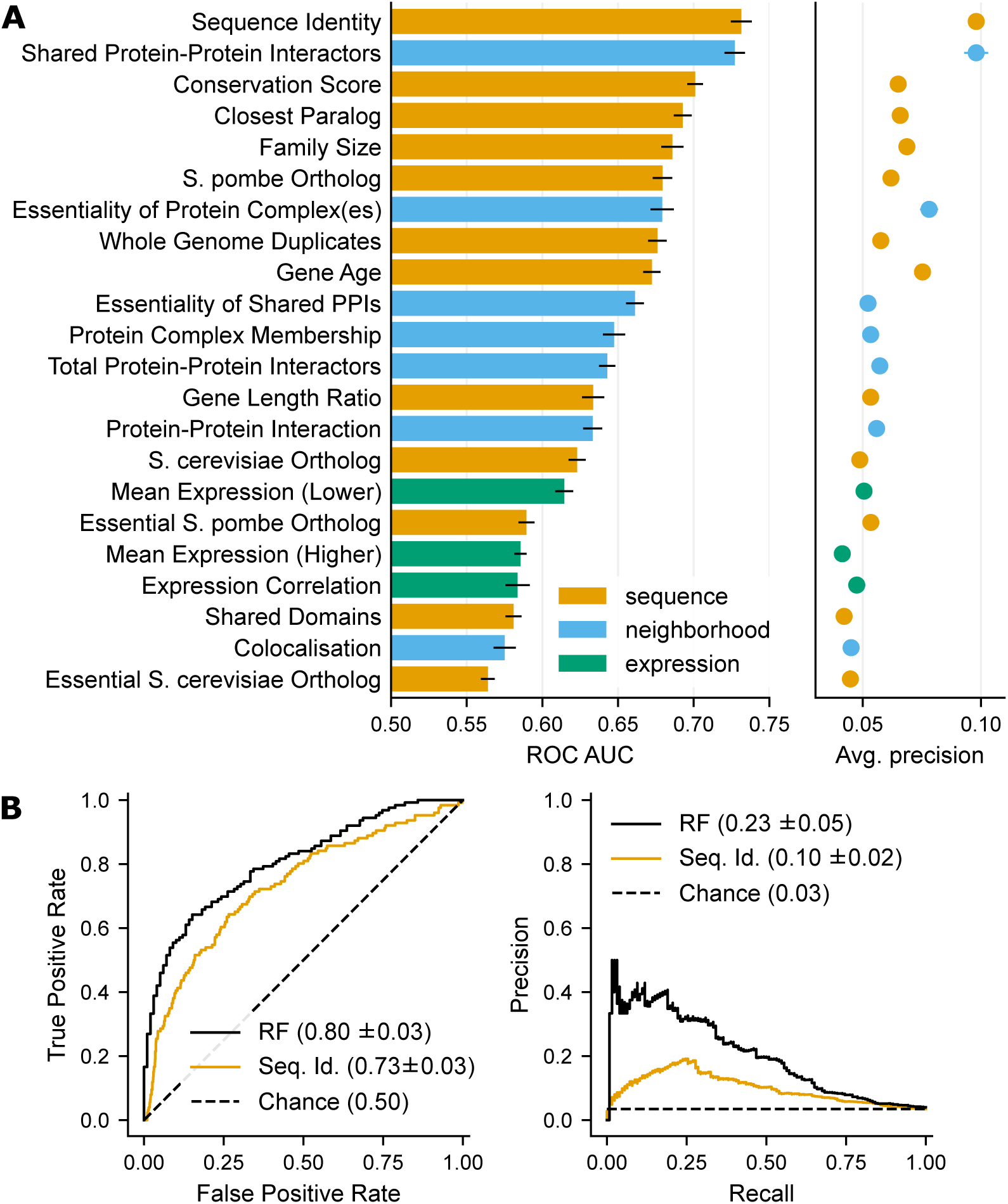
Assessing the predictive power of 22 features of paralog pairs and a random forest classifier. (A) Bar chart showing the area under the ROC curve (ROC AUC) and point plot showing the average precision for each feature as an individual classifier over our dataset of SL and non- SL paralog pairs. The error bars indicate the mean standard error based on computing the ROC AUC and average precision for 10 randomly selected, stratified fourths of the dataset. Bars and dots are colored according to the feature category. (B) ROC and PR curves for the random forest classifier (RF, black), based on stratified 4-fold cross-validation of our dataset, and for sequence identity alone (orange). The values shown in the legend are the ROC AUC and average precision, along with standard deviation where applicable.

The two individual features that best predict synthetic lethality in our dataset are amino acid sequence identity and shared protein-protein interactors (measured as the -log10 *p*-value from a Fisher’s exact test of the overlap of A1 and A2’s interactors) (Fig. 3A). These features both have a ROC AUC of ∼0.73 and an average precision of ∼10%. Both features can be considered measurements of ‘functional similarity’ - with sequence identity measuring how similar the paralogs’ protein sequences are and shared protein-protein interactions measuring how similar the two paralogs are in terms of their role in the protein-protein interaction network. Other features that measure similarity (fraction of shared domains, co-localisation, and co-expression) are predictive of SL but to a much lesser extent (Fig. 3A).

Features that capture evolutionary conservation also appear to be highly predictive of synthetic lethality, in particular a measure of the number of species in which a paralog pair has an identifiable ortholog (‘Conservation Score’, see Methods). Genes that have been conserved over a long evolutionary period are more likely to encode essential functionality (Chen et al., 2012), which could explain why highly conserved paralog pairs are more likely to be SL. Similarly, we found that gene age (Capra et al., 2012), a measure of the phylogenetic age of a protein, is very predictive of SL.

We included a number of features relating to gene conservation in the model eukaryotes *S. cerevisiae* and *Schizosaccharomyces pombe* (Fig. 3A, Table S6 and Methods for feature details). Although both are yeasts, they are very evolutionary distant (over 400 million years (Sipiczki, 2000)) and differ significantly in terms of their basic biology (e.g. *S. pombe* has RNAi machinery and splicing much more similar to metazoans), and in terms of their genome composition (*S. cerevisiae* has undergone a whole genome duplication while *S. pombe* has not (Wood, 2006)). For both species we have a comprehensive overview of which genes in their genome are essential, thanks to large scale gene deletion projects (Giaever et al., 2002; Kim et al., 2010). We anticipated that having an essential *S. cerevisiae* or *S. pombe* ortholog would be very predictive of an SL relationship between a human paralog pair, but surprisingly we found that these two features were only weakly predictive. In fact, having any identifiable ortholog in *S. pombe* was more predictive of SL than having an essential *S. pombe* ortholog (average precision ∼6.2% vs. ∼5.3%), and the same trend was evident for *S. cerevisiae* (average precision ∼4.9% vs. ∼4.5%) (Fig. 3A). One possible explanation for this is that the orthologs in the two yeast species may themselves have been subject to a gene duplication event and hence may appear non-essential due to paralog buffering. However, even when restricting our analysis to cases in which there was a single ortholog in either species (i.e. a many-to-one mapping) the addition of essentiality information did not improve predictive power (Fig. S2). This suggests that in general having any identifiable ortholog in budding or fission yeast is more informative than having an essential ortholog.

Species conservation and protein age are both indicators of gene importance – as noted, a gene that is widely conserved is likely to perform some essential function. We also included several more direct measures of gene importance derived from gene essentiality profiles calculated from CRISPR screens in cancer cell lines. We anticipated that paralog pairs that function in largely essential protein complexes (i.e. complexes with subunits that are frequently essential across the DepMap cell lines) or that have protein-protein interactions with essential genes are more likely to be SL and found that this is the case (the average precision associated with protein complex and shared PPI essentiality is ∼7.8% and ∼5.2% respectively) (Fig. 3A).

Previous work has suggested that genes that encode proteins with a high number of PPIs are more likely to be essential, although the reasons for this association are much disputed (Hahn and Kern, 2005; He and Zhang, 2006; Wang et al., 2009; Zotenko et al., 2008). We tested whether paralog pairs with more PPIs are more likely to be SL and found that this is indeed the case (Fig. 3A, ROC AUC ∼0.64, average precision ∼6%). However, this feature was nowhere near as predictive as shared PPIs (ROC AUC ∼0.73, average precision ∼10%), suggesting that the total number of interactions that the genes in a paralog pair have is less important than the fraction they have in common.

The final set of features we found to be predictive relate to the possibility of compensation by other paralogs in a family. In many cases a pair of paralogs belongs to a much larger family of paralogs. We found that SL paralogs tend to come from smaller families (the median family size for SL paralogs is 5, compared to 8 for non-SL paralogs) and are more likely to be each other’s closest paralog (Fig. 3A).

In addition to evaluating the predictive power of different features using ROC AUC and average precision, we used standard statistical approaches to determine, for all features, whether SL pairs have significantly different distributions than non-SL pairs (Fig. S3, Table S6). For all quantitative features we find that SL pairs have significantly different feature values than non-SL pairs (all Mann-Whitney test *p*<0.001); e.g. the mean essentiality of shared PPIs is significantly higher for SL pairs than for non-SL pairs (*p*=1.8x10^-10^). Similarly, for all Boolean features we find that SL pairs are significantly enriched for specific feature values (all Fisher’s exact test *p*<0.001); e.g. SL pairs are significantly more likely to have an *S. pombe* ortholog than non-SL pairs (*p*=2.1x10^-16^).

Overall, the features we have assembled for paralog pairs quantify 1) the extent of their functional similarity, e.g. sequence identity or co-expression; 2) whether their shared functionality is likely to be essential, e.g. membership of an essential protein complex; and 3) whether there are other potential sources of genetic redundancy that would mask a synthetic lethal relationship, e.g. paralog family size. We observe that paralog pairs are likely to be SL if they are functionally similar, if their shared function is important, and if they can uniquely compensate for each other’s loss.

### An ensemble classifier to predict synthetic lethality among paralog pairs

Having identified several features that are individually predictive of synthetic lethality between paralog pairs in our dataset, we next asked whether combining these features into an ensemble classifier could provide greater predictive power. Though several of the features are expected to be correlated with each other, we retain all features as they can still contribute unique information to the classifier. For example, we previously found that whole genome duplicates are more likely to be in protein complexes, and while this could partially explain why whole genome duplicates are more likely to be synthetic lethal, we also found that whole genome duplication was still predictive when we controlled for protein complex membership (De Kegel and Ryan, 2019).

We trained a random forest classifier (Breiman, 2001), i.e. an ensemble of decision trees, using our dataset of SL and non-SL paralog pairs annotated with all 22 features (see Methods). A tree-based ensemble classifier makes it possible to capture interactions between features as well as non-linear relationships between a given feature and paralog synthetic lethality. We find that the predictions made by the random forest classifier greatly outperform sequence identity, one of the best individual features, which we use as the baseline for comparison (Fig. 3B). Specifically, the classifier achieves a ROC AUC of ∼0.80 and average precision of ∼23% over the training data, while the metrics for sequence identity were ∼0.73 and ∼10% respectively.

### An ensemble classifier can accurately predict synthetic lethality in combinatorial CRISPR screens

We found that our classifier markedly outperformed sequence identity alone at distinguishing SL from non-SL paralog pairs in our training dataset. To evaluate the classifier using an orthogonal dataset, we next evaluated its ability to predict the results of the four published combinatorial CRISPR screens of paralog pairs (Dede et al., 2020; Gonatopoulos-Pournatzis et al., 2020; Parrish et al., 2020; Thompson et al., 2021). Given that many of the hits in these screens are cell line and/or screen specific (Fig. 1E-F), but we are mainly interested in identifying robust SLs, we first evaluated our classifier on the consensus results from all four screens. In total 1,845 of the Ensembl reference paralog pairs were tested in at least one CRISPR screen while 545 pairs were tested in at least two screens (this excludes Thompson *et al*. pairs that were filtered out due to single gene essentiality, see Methods). We used these 545 pairs to form a screen consensus dataset – treating pairs that were identified as SL in at least two screens as true positives (n=50) and those that were screened in at least two screens but not identified as SL in any screen as true negatives (n=407). We find that our classifier can make accurate predictions for this screen consensus dataset – up to a recall of 32% we observe precision of at least 50% (Fig. 4A). In addition, we note that our classifier clearly outperforms sequence identity – the average precision for our predictions is ∼49%, compared to ∼20% for sequence identity alone. By chance the average precision for this dataset is the hit rate, i.e. ∼11%, which is similar to the hit rate and by extension the class imbalance that we observed for the individual screens (Fig. 1E). We repeated this analysis for hits identified in one or more screens (n=224) and found that our classifier again outperforms sequence identity alone (average precision 38% vs 12% for sequence identity, Fig. S4A).

**Figure 4.**
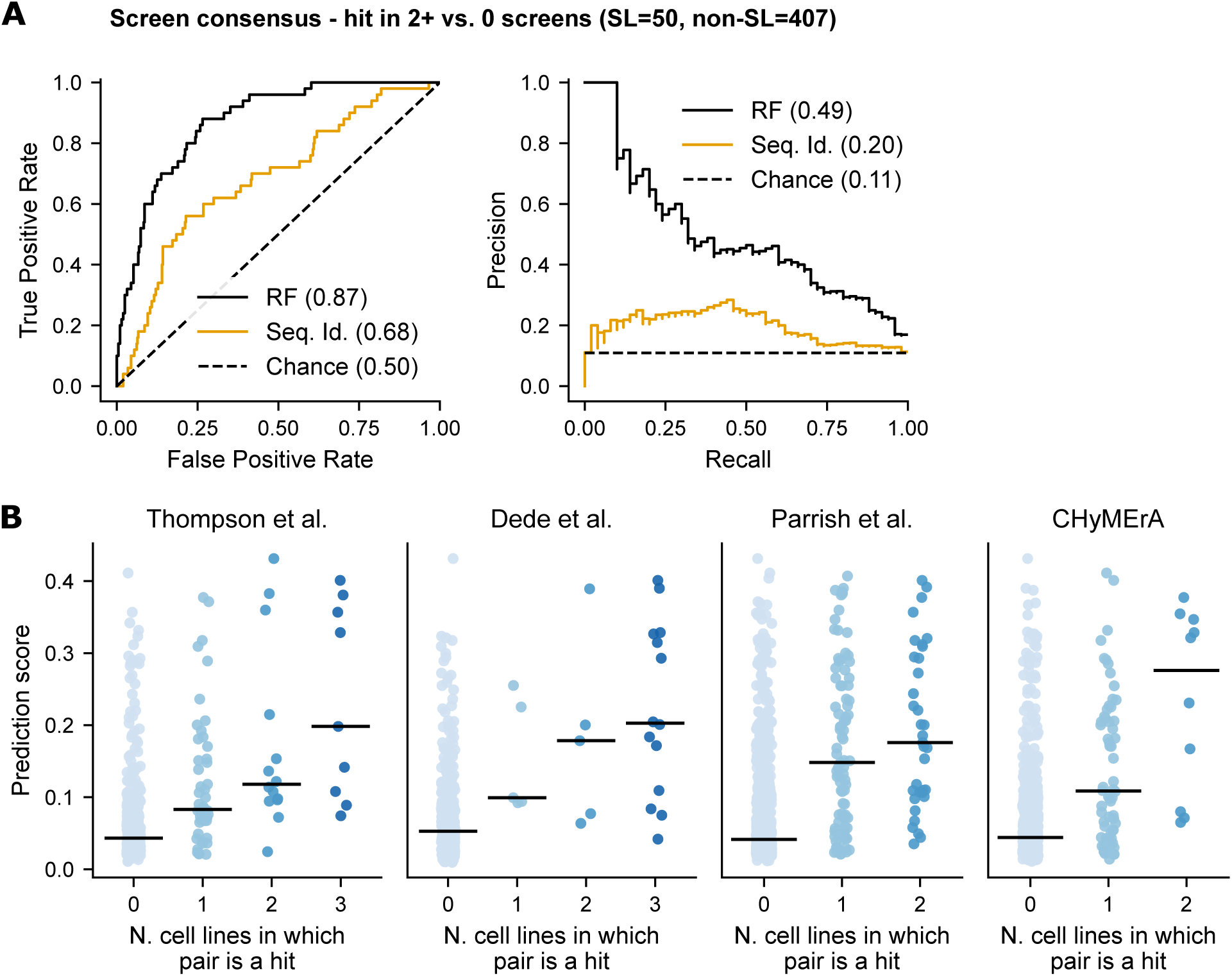
Assessing the performance of our classifier on data from independent genetic interaction screens. (A) ROC and PR curves showing the performance of our predictions (black) and sequence identity (orange) in distinguishing screen consensus SLs (pairs SL in 2+ combinatorial screens) from non-SLs (pairs tested in 2+ screens and not found to be SL in any screen). The values shown in the legends are ROC AUC and average precision. (B) Strip plots showing the individual prediction scores for pairs identified as SL in 0, 1, 2 or 3 cell lines in each combinatorial screen. The black lines indicate the median prediction score for each category (number of cell lines).

Some of the paralog pairs tested in the combinatorial screens were also present in our training dataset, so to ensure that our classifier wasn’t overfitting to the training data we repeated the evaluation after removing pairs present in our dataset (as either an SL or non-SL pair). We found that our classifier still performs well when only considering these unseen pairs (average precision ∼46% for consensus hits, Fig. S4B, and ∼32% for hits in any screen, Fig. S4C) indicating that the classifier’s performance on the results of the combinatorial screens is not just based on overfitting to the common pairs.

Although we are primarily interested in the classifier’s performance on the screen consensus results, we also evaluated our classifier on each of the four combinatorial screens independently (Fig. S5). For all screens we found that our classifier outperforms sequence identity both at identifying robust SLs (hits in two or more cell lines) and identifying all SLs (hits in any cell line screened).

There are no previous methods for specifically predicting SL between paralog pairs, but there are several approaches for predicting SL between gene pairs in general (i.e. between both paralog and unrelated gene pairs). To further evaluate our classifier, we compared its performance to three previously developed SL classifiers: the first uses PPI network topology patterns and is trained on *S. cerevisiae* SL data (SINaTRA) (Jacunski et al., 2015); the second uses a wider set of conserved PPI network topology patterns and is trained on SL data from humans and four other species (SLant) (Benstead-Hume et al., 2019); and the third attempts to predict new edges in the SL interaction network based solely on the existing network structure (Cai et al., 2020). Due to the input requirements of each of these classifiers, not all screened paralog pairs could be assigned a predicted score; e.g. the Cai *et al*. classifier can only make predictions for pairs where both genes are already present in the training SL network. Consequently we focussed our comparison on those pairs for which predictions could be made. Our classifier outperforms the SINaTRA, SLant and Cai et al. classifiers at identifying screen consensus SLs (hits in two or more screens) (Fig. S6) and all SLs (hits in any screen), including when only considering the paralog pairs that were not in our training dataset (unseen pairs) (Table S7). This suggests that in comparison to a generic SL classifier, our paralog-specific SL classifier is better able to capture the factors of a potential synthetic lethal relationship between paralog pairs. In general it should be noted that many existing generic SL classifiers can only make predictions for a subset of all ∼36.6k paralog pairs. Despite making predictions for millions of gene pairs the SINaTRA, SLant and Cai *et al*. classifiers respectively only provide predictions for ∼56.5% (n=20,701), ∼9.2% (n=3,385) and ∼10% (n=3,672) of all paralog pairs.

We note that in addition to methods for predicting synthetic lethality in humans, there are also methods for prioritising experimentally determined SLs that might be of greatest clinical interest, e.g. ISLE (Lee et al., 2018). However these approaches require experimentally determined SLs as input and consequently cannot be used to prioritise most paralog pairs (e.g. ISLE makes predictions for only 133 paralog pairs).

### Distinguishing robust synthetic lethal interactions from cell-line specific synthetic lethals

To assess whether our classifier could distinguish robust synthetic lethal interactions, observed in multiple cell lines, from those seen in only a single cell line we made predictions for the full set of paralog pairs tested by each combinatorial CRISPR screen (Dede et al., 2020; Gonatopoulos-Pournatzis et al., 2020; Parrish et al., 2020; Thompson et al., 2021). We split each dataset according to the number cell lines in which the pair was identified as SL and found that our classifier captures some information as to which SL relationships might be more robust; on average, pairs that were hits in three cell lines have a higher predicted score than those that were hits in only two, which in turn have higher scores than those that were hits in only one cell line (Fig. 4B). This trend is clear for all four screens and comparing the prediction scores for robust vs. cell-line specific hits we find that they are significantly different in two cases (two-sided Mann-Whitney test: Thompson *p*=0.005, Dede *p*=0.57, Parrish *p*=0.18, CHyMErA *p*=0.02). We also note that in all cases there is a significant difference in the prediction scores for hits in zero vs. one cell line (two-sided Mann-Whitney test, *p*<0.01).

### Understanding why specific paralog pairs are likely or unlikely to be synthetic lethal

Having validated our classifier using data from four combinatorial CRISPR screens, we next made predictions for all ∼36.6k paralog pairs from Ensembl that have at least 20% sequence identity (in at least 1 direction of gene-to-gene comparison) (Table S8). These scores provide a means of ranking all paralog pairs according to their likelihood of being synthetic lethal. We found that 11/12 SLs with low throughput experimental validation (Table S1) and 47/50 pairs from the screen consensus rank in the top 5% of our predictions (Fig. 5A).

**Figure 5.**
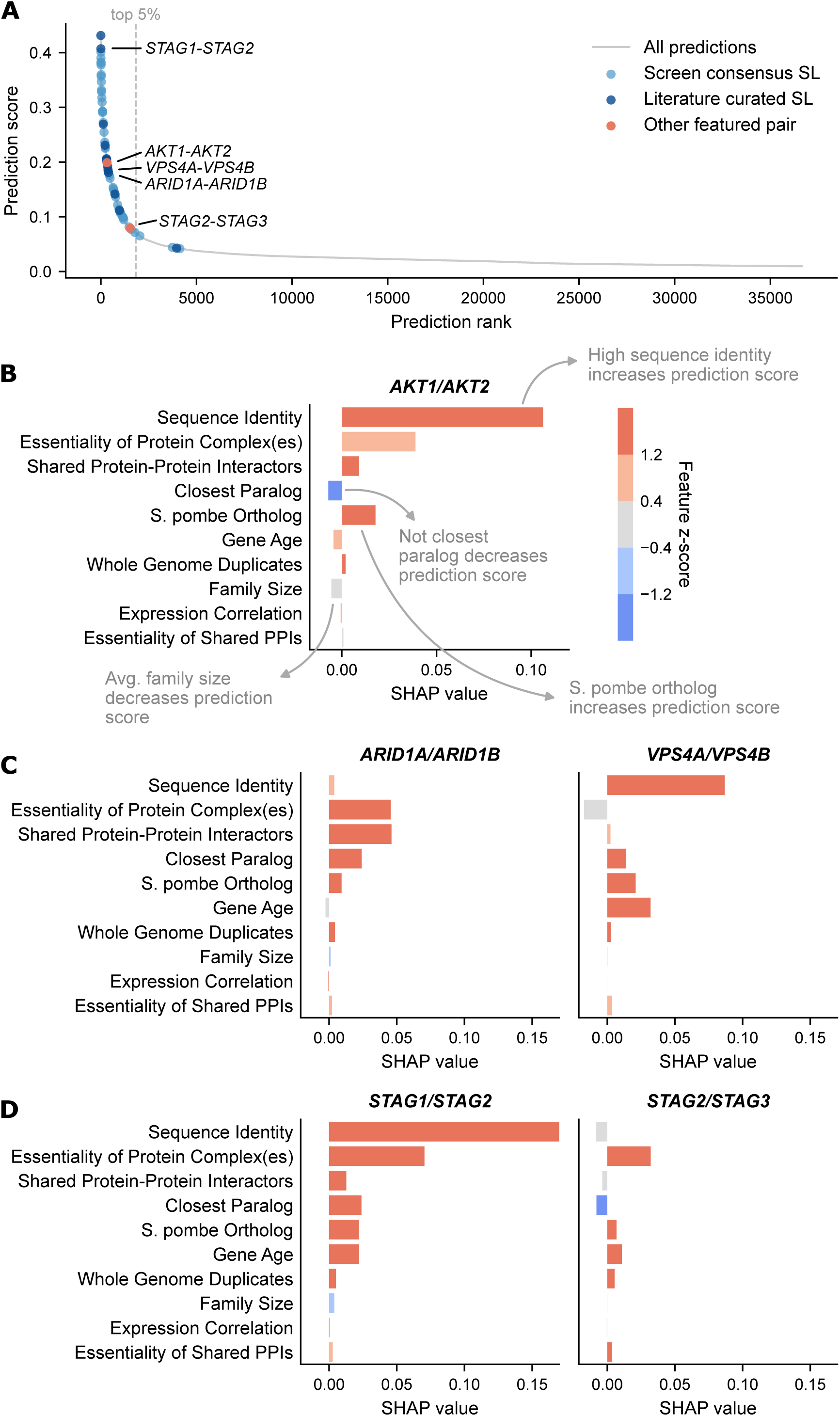
Predictions for all paralog pairs and some of their local explanations. (A) Plot showing the prediction score vs. rank for all paralog pairs. The light blue dots are predictions for screen consensus SLs (pairs SL in 2+ combinatorial screens), while the dark blue dots are predictions for pairs that have been previously validated as SL (Table S1). The orange dots are predictions for other pairs featured in B-D. All featured pairs are annotated. Predictions left of the dotted grey line are in the top 5%. (B) Annotated SHAP profile for the pair *AKT1/AKT2.* SHAP values are shown on the x-axis and can be either positive (feature increases prediction score) or negative (feature decreases prediction score). The 10 features with the largest global impact are shown on the y-axis; this feature subset was determined by calculating the median absolute SHAP value associated with each feature across all paralog pairs. The hue indicates the feature z-score, i.e. whether the feature has a relatively high (dark orange) or relatively low (dark blue) value for the given pair. (C) SHAP profiles for the pairs *ARID1A/ARID1B* and *VPS4A/VPS4B*. (D) SHAP profiles for the pairs *STAG1/STAG2* and *STAG2/STAG3*.

Although we identified a number of features that are individually predictive of SL (Fig. 3A), we noted that not all of the pairs with high predicted scores have what might be expected values for these features. For example, in general pairs with high-sequence identity are more likely to be SL, but a number of highly ranked pairs have only moderate sequence identity (e.g. *UBB*/*UBC*, a pair which is not present in our dataset but which has previously been validated as SL (Table S1) (Tsherniak et al., 2017), is ranked in the top 3% of gene pairs but has sequence identity close to the average of all paralog pairs). Similarly, in general pairs from small paralog families are more likely to be SL but some paralog pairs from larger families are highly ranked (e.g. *RAB1A/RAB1B*, a pair not present in our dataset but recently identified as SL in a high-throughput screen (Blomen et al., 2015), is ranked in the top 1% of pairs but belongs to a paralog family of size 15). To understand which factors influence the predictions for a given paralog pair, we made use of TreeExplainer, a recently developed algorithm to compute local explanations for ensemble tree-based predictions (Lundberg et al., 2020). Briefly, TreeExplainer computes how much each feature pushes a specific prediction higher or lower. This numeric measure of feature credit is termed a SHAP (SHapley Additive exPlanations) value. More technically speaking, the SHAP values for a given paralog pair sum to the difference between the prediction for that pair and the average prediction for the classifier (i.e. the average prediction over all pairs in the training data). To complement the SHAP values (Table S9) we also computed z-scores for all the feature values (see Methods); this provides us with a SHAP profile that links feature values and feature impacts (on prediction) for each pair. An example SHAP profile, for *AKT1/AKT2*, is shown in Fig. 5B.

The SHAP profiles provide a visual overview of what features influence the prediction for a specific paralog pair. For example, the paralog pairs *ARID1A/ARID1B* and *VPS4A/VPS4B* are similarly ranked and have both been previously validated as SL (Helming et al., 2014; Lord et al., 2020; Szymańska et al., 2020) (Table S1). A comparison of their SHAP profiles reveals that the two pairs are highly ranked for very different reasons (Fig. 5C). The profile for *ARID1A/ARID1B* shows that its prediction is mainly influenced by its membership of essential protein complex(es) and a significant overlap in the protein-protein interactors of *ARID1A* and *ARID1B*. The sequence identity of the two proteins, which is moderate, does not significantly influence the prediction. In contrast, the prediction for *VPS4A/VPS4B* is mainly driven by their high sequence identity and old evolutionary age.

In cases where a gene has multiple paralogs, the SHAP profiles can be used to explain why one paralog pair is ranked more highly than others. For example, the tumour suppressor *STAG2* has two paralogs, *STAG1* and *STAG3*, but *STAG1/STAG2* is ranked significantly higher (in fact it is one of the highest ranked pairs) than *STAG2/STAG3* (Fig. 5A). Consistent with these rankings *STAG1*, but not *STAG3*, was recently identified as a hit in a genome-wide CRISPR screen for *STAG2* synthetic lethal interactions (van der Lelij et al., 2020). Looking at the SHAP profile for *STAG1/STAG2,* we observe that its prediction was pushed up substantially by high sequence identity and membership in essential complex(es), and to a lesser extent by some other features, such as shared protein-protein interactors and *STAG1* and *STAG2* being each other’s closest paralogs (Fig. 5D, left). While still being pushed up by its membership of essential complex(es), the prediction for *STAG2/STAG3* is simultaneously pushed down by its moderate sequence identity, lack of shared interactors, and the genes not being each other’s closest paralogs (Fig. 5D, right). This example indicates that SHAP profiles can reveal why one paralog pair from a paralog family is ranked higher than another despite the two pairs having some common characteristics. Fig. S7 provides an additional example, showing why *AKT1* is predicted to be SL with *AKT2* and *AKT3* but not with its 11 other paralogs; *AKT1/AKT2 and AKT1/AKT3* were also previously identified as SL in a combinatorial screen (Najm et al., 2018).

### *ASF1A/ASF1B* and *COPS7A/COPS7B* are synthetic lethal pairs

We selected two of our highly ranked predicted synthetic lethal interactions for further validation: *ASF1A/ASF1B* and *COPS7A/COPS7B* (Fig. 6A). Neither pair is present in our computational dataset of SLs nor in our manually curated set of validated SLs (Table S1). We selected *ASF1A/ASF1B* for validation because it is among the highest ranked predictions (top 0.1%) and because *ASF1A* was previously reported as frequently homozygous deleted in multiple cancer types including prostate adenocarcinoma and diffuse large B cell lymphoma (Lee et al., 2017). Re-analysis of the Cancer Genome Atlas confirms these two previous associations and suggests that ASF1A may also be recurrently deleted in uveal melanoma (Fig. 6B) (Cerami et al., 2012; Gao et al., 2013; Hoadley et al., 2018; Robertson et al., 2017). The SHAP profile for *ASF1A/ASF1B* reveals that it is highly ranked among our predictions because the two paralogs share high sequence identity (70%), have a high proportion of shared interactions, and function as part of multiple largely essential complexes (Histone H3.1 and H3.3 complexes (Tagami et al., 2004)) (Fig. 6C). To test this synthetic lethality we used the same approach as in (Lord et al., 2020) – we obtained knockouts of *ASF1A* and *ASF1B* in the near haploid HAP1 cell line and tested whether these knockout cells were more sensitive to the inhibition, via siRNA, of the other paralog. In both knockouts we found this to be the case – *ASF1A* knockout cell lines were more sensitive than wild-type parental cells to the inhibition of *ASF1B* while *ASF1B* knockouts were more sensitive to the inhibition of *ASF1A* (Fig. 6D, Table S10). The fitness impact of the siRNA targeting *ASF1A/ASF1B* in the wild-type cells was similar to that observed for a scrambled siRNA control, suggesting that inhibition of either gene in wild-type cells is well tolerated.

**Figure 6.**
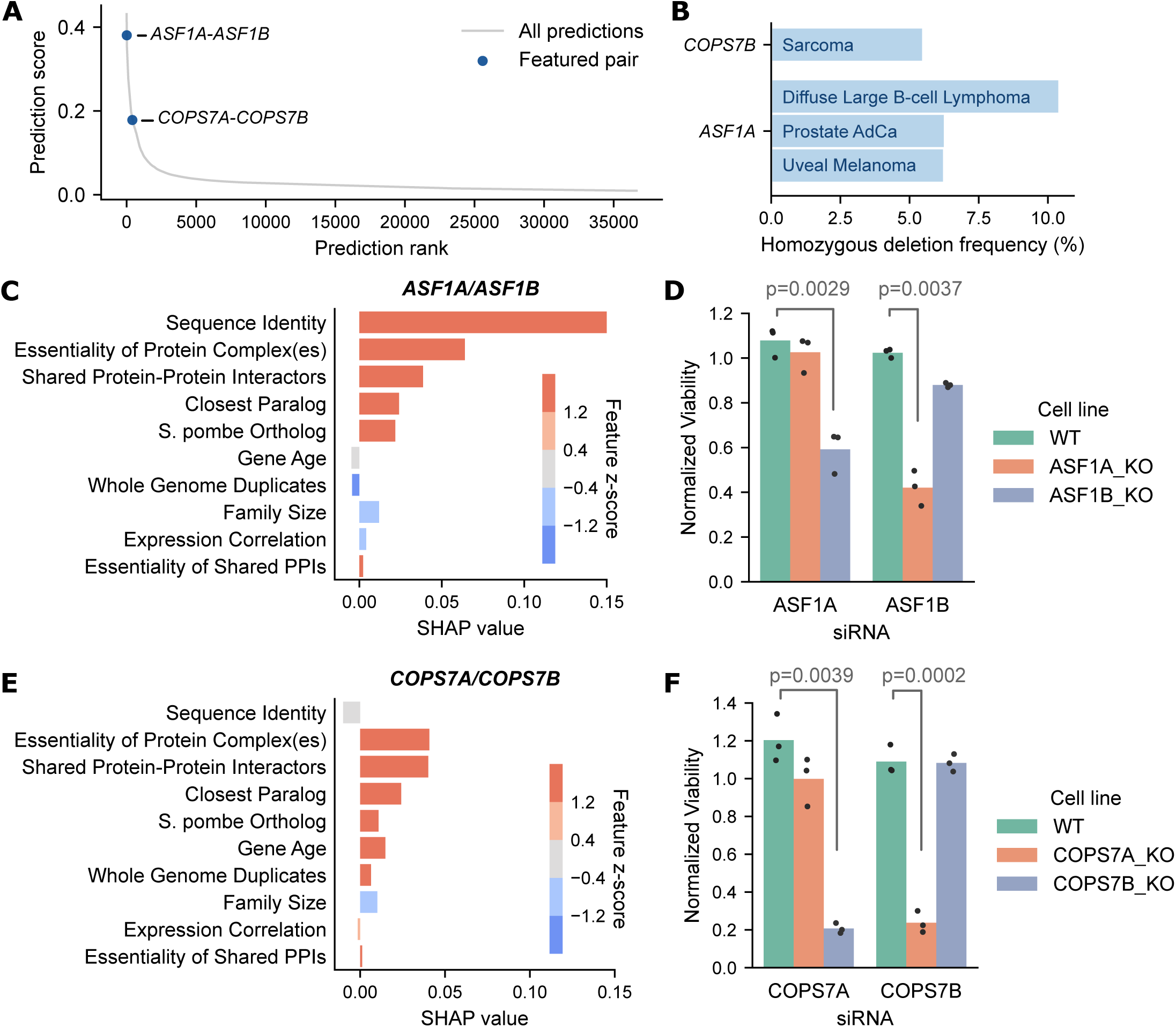
Prediction and validation of two new paralog synthetic lethalities. (A) Plot showing the prediction score vs. rank for all paralog pairs (as in Fig. 5A), with the two pairs featured in B-F highlighted in dark blue. (B) Bar chart showing the homozygous deletion frequency of *COPS7B* and *ASF1A* in the indicated cancer types, based on tumour samples from the TCGA (AdCa=Adenocarcinoma). (C) SHAP profile for the paralog pair *ASF1A/ASF1B*. (D) Mean viability of HAP1 cells treated with siRNA smartpools targeting *ASF1A* or *ASF1B*. Individual data points are shown as black dots. Data are normalized within each cell line such that the mean viability of cells treated with a negative control (non-targeting scrambled siRNA) is equal to one and the mean viability treated with a positive control (siRNA smartpool targeting the broadly essential PLK1 gene) is equal to 0. The p-values come from two sided heteroscedastic t-tests. (E) SHAP profile for *COPS7A/COPS7B*. (F) Bar chart showing the mean viability of HAP1 cells treated with siRNA smartpools targeting *COPS7A* or *COPS7B*; normalization and statistics as in (D).

We next sought to validate the synthetic lethality between *COPS7A* and *COPS7B.* Through analysis of tumour copy number profiles from the Cancer Genome Atlas, we found that *COPS7B* is subject to frequent homozygous deletion in sarcoma (Fig. 6B) (Cancer Genome Atlas Research Network, 2017; Cerami et al., 2012; Gao et al., 2013; Hoadley et al., 2018). *COPS7A/COPS7B* is further down the list of predictions (Fig. 6A, top 1.5%) than *ASF1A*/*ASF1B* and has significantly lower sequence identity (47%). The SHAP profile reveals that the prediction for this pair is pushed down slightly by the moderate sequence identity but pushed up by a high proportion of shared protein-protein interactions and membership of an essential complex (the COP9 Signalosome (Seeger et al., 1998)) (Fig. 6E). We used the same approach to test the synthetic lethality between this pair and again found that *COPS7A* knockout cells were more sensitive to the inhibition of *COPS7B* and vice-versa (Fig. 6F, Table S9).

## Discussion

In this work we have developed an approach for predicting synthetic lethality among human paralog pairs. We established a dataset of (non-)SL paralog pairs that allowed us to evaluate the predictive power of 22 diverse features of paralog pairs, as well as a random forest classifier that incorporates these features. We found that sequence identity and shared protein-protein interactors are the best individual predictors of synthetic lethality between paralogs. However, the predictions made by our classifier significantly outperform these features alone. Furthermore, we show that our classifier, trained with data from single gene perturbation screens, can accurately predict the consensus results from four independent combinatorial CRISPR screens of paralog pairs (Dede et al., 2020; Gonatopoulos-Pournatzis et al., 2020; Parrish et al., 2020; Thompson et al., 2021). Finally, we generated synthetic lethality predictions, with local explanations based on feature credit, for ∼36.6k paralog pairs. These predictions and their explanations can be viewed at cancergd.org/static/paralogs.

### A computational dataset of robust synthetic lethal paralog pairs

We have developed a dataset of 126 robust synthetic lethal and 3,508 non-synthetic lethal paralog pairs. While this dataset likely contains both false positives (pairs that are incorrectly identified as SL) and false negatives (pairs that are incorrectly identified as not SL), we anticipate that there are fewer false positives as we have selected a relatively stringent false discovery rate (10%) and controlled for factors that might contribute to false positives. In particular we excluded gene pairs with tissue specific expression patterns and incorporated lineage as a covariate into our analysis to identify synthetic lethality. This should prevent us calling a synthetic lethal relationship in cases where paralogs might simply be essential in a specific lineage because they are only expressed in that lineage. We anticipate that there are a significantly higher number of false negatives – the principal reason for this is that our ability to identify genes with loss-of-function alterations is imperfect. We may have called gene loss where a gene only has a reduction in expression or copy number, or a heterozygous loss-of-function mutation. Such limitations could be overcome by, e.g. incorporating protein expression datasets for the same cell lines (Nusinow et al., 2020), but currently this would severely limit the number of cell lines and paralog pairs we could analyse. Despite its limitations we show that our dataset can be used to build a classifier that can accurately predict synthetic lethality in combinatorial screens, suggesting that it is more than sufficient to learn general trends that distinguish SL from non-SL paralog pairs. We anticipate that as more combinatorial screens of paralog pairs are performed it may be possible to improve our dataset by integrating these with our analysis of the DepMap data, e.g. only calling positives/negatives that are supported by both types of screen. Due to the bias of the extant screens doing so currently would filter out the majority of paralog pairs from our dataset.

### Frequency of synthetic lethality between paralogs in human cell lines

Our dataset of human paralog (non-)SLs has a synthetic lethality rate of ∼3.5%, which is significantly lower than that reported for budding yeast, where 25-35% of paralog pairs have been identified as SL. However, this lower frequency of synthetic lethality is reasonably consistent with the results of the four combinatorial screens of paralog pairs analysed in this study. Thompson *et al*. (Thompson et al., 2021), Dede *et al*. (Dede et al., 2020), Parrish *et al*. (Parrish et al., 2020) and Gonatopoulos- Pournatzis *et al*. (Gonatopoulos-Pournatzis et al., 2020) performed combinatorial CRISPR screens in 2-3 cell lines to identify synthetic lethality between paralog pairs. All four screens had significant selection bias in terms of the paralogs screened (Fig. 1A-D) which might be expected to increase the synthetic lethality rate. However, all screens still had a relatively low synthetic lethality rate: only ∼4.8% (Thompson et al., 2021), ∼4.8% (Dede et al., 2020), ∼3.4% (Parrish et al., 2020) and ∼1.5% ((Gonatopoulos-Pournatzis et al., 2020) filtered to SL pairs) of screened paralog pairs were identified as hits in two or more cell lines. One explanation for this reduced SL rate in human cells is that the median paralog family size is larger in humans (4 genes) than in *S. cerevisiae* (2 genes), in line with the ancestor of humans having undergone an extra round of whole genome duplication (Dehal and Boore, 2005). The consequences of perturbing any individual paralog pair in human cells may thus be masked by the presence of additional paralogs. Such higher-order relationships could be revealed by perturbing three or more genes simultaneously, as has been done in budding yeast (Haber et al., 2013; Kuzmin et al., 2020).

### What makes a paralog pair likely to be synthetic lethal?

In this work we tried to answer this question generally, through analysis of the predictive power of individual features of paralog pairs (Fig. 3A) and on a pair-specific level, through analysis of the SHAP profiles of our predictions (Fig. 5-6).

Overall, we found that synthetic lethal paralog pairs are typically older and more widely conserved than non-synthetic lethal pairs. They are on average more highly expressed and are more likely to interact with essential genes. These findings suggest that the general importance of the genes’ function is a key factor in making a paralog pair SL. Another key factor seems to be the degree of functional similarity between the genes – we found that SL paralog pairs tend to have higher sequence identity and greater overlap of protein-protein interactors compared to non-SL pairs. Finally we found that although many paralogs belong to large families, SL pairs generally come from smaller families, suggesting that a paralog pair is more likely to be SL if the two genes uniquely compensate for each other’s loss.

Interestingly we found that a specific paralog pair can be highly ranked by our classifier even if it doesn’t follow the overall trends that we observed. For example, although SL paralog pairs generally have higher sequence identity compared to non-SL paralog pairs, we identified several pairs (e.g. *ARID1A/ARID1B* and *COPS7A/COPS7B*) that are highly ranked despite average or below average sequence identity. This suggests that the causes of SL among paralog pairs can differ significantly, and highlights the importance of using a set of paralog pairs without prior feature bias to analyse the factors that make a paralog pair likely to be SL.

### Robust vs. cell-line specific synthetic lethal interactions

The SL pairs in our dataset were identified from combined analysis of heterogeneous cancer cell lines and are thus expected to be largely robust to cancer type and genetic background. There are likely additional context-specific synthetic lethal relationships (Henkel et al., 2019; Ryan et al., 2018) among our candidate paralog pairs but we cannot detect them here. In contrast, the paralog pairs screened in combinatorial screens (Dede et al., 2020; Gonatopoulos-Pournatzis et al., 2020; Parrish et al., 2020; Thompson et al., 2021) are initially called SL in a cell-line specific fashion. We found that on average our classifier assigned higher prediction scores to pairs that were identified as SL in multiple cell lines, compared to pairs that were identified as SL in only one cell line. This suggests that our classifier, perhaps due to the training data, has some ability to distinguish robust synthetic lethalities from those that might be cell-line specific. Our classifier is based on context- agnostic features, e.g. average rather than cell-line specific complex essentiality, and can thus only make context-agnostic predictions of SL. However, as genetic interactions screens are carried out across a wider variety of cell lines it may become possible to develop a classifier that can make context-specific predictions.

### *ASF1A/ASF1B* and *COPS7A/COPS7B* are validated synthetic lethal pairs

We have demonstrated, using a low-throughput approach, that *ASF1A*/*ASF1B* and *COPS7A/COPS7B* are synthetic lethal in the HAP1 cell line. These paralog pairs represent a very high ranking prediction (*ASF1A/ASF1B*, top 0.1%) and a lower ranking prediction (*COPS7A/COPS7B,* top 1.5%) and a pair with high sequence identity (*ASF1A/ASF1B*, 70% id) and moderate sequence identity (*COPS7A/COPS7B*, 47% id). We note that *ASF1A*/*ASF1B* was also identified as a robust hit in two CRISPR-Cas9 screens published while this manuscript was in preparation (Parrish et al., 2020; Thompson et al., 2021), while *COPS7A/COPS7B* was screened by both Thompson *et al*. and Dede *et al*. (Dede et al., 2020), but only identified as a hit in Dede *et al*. Further work is required to establish if this is due to context-specific effects or if it is a false-negative in the Thompson *et al*. screen. Due to the relatively high frequency of homozygous deletion of *ASF1A* in different cancer types (Lee et al., 2017) and *COPS7B* in sarcoma (Fig. 6B), both synthetic lethal interactions may be of use as therapeutic targets.

### Ranking paralog pairs most likely to have a synthetic lethal relationship

Paralogs are a rich source of synthetic lethality and exploiting this could be a widely applicable approach to targeting gene loss in cancer. However, there is a large number of paralog pairs in the human genome (>36k), and only a small fraction have been tested to date. The predictions made by our classifier provide a ranking of all paralog pairs in terms of how likely they are to exhibit a robust synthetic lethal relationship. In contrast, many existing genome-wide approaches to predicting SL only make predictions for a fraction of all paralog pairs. In addition, we have demonstrated that our classifier significantly outperforms three previously developed SL classifiers (Benstead-Hume et al., 2019; Cai et al., 2020; Jacunski et al., 2015) in distinguishing paralog pairs that were identified as (robust) hits in combinatorial CRISPR screens. Our rankings can serve as a resource for selecting paralog pairs for inclusion in future combinatorial screens. To facilitate such analyses we have provided full details of our predictions, along with their SHAP profiles, in the supplementary materials. We have also made these easy to explore using the supplementary site cancergd.org/static/paralogs

## Methods

Unless otherwise stated, all computational analysis was carried out using Python 3.7, Pandas 1.0.1 (McKinney, 2011), SciPy 1.4.1 (Virtanen et al., 2020), StatsModels 0.11.0 (Seabold and Perktold, 2010), numpy 1.18.5 (Harris et al., 2020) and scikit-learn 0.23.1 (Pedregosa et al., 2011). Many of the figures were created with Seaborn 0.11.1 (Waskom, 2021). Notebooks containing the code for this project are available at https://github.com/cancergenetics/paralog_SL_prediction

### Paralog pairs reference list

Protein-coding paralog gene pairs and their amino acid sequence identities were obtained from Ensembl 93 (GRCh38.p12 genome assembly) (Zerbino et al., 2018). These pairs were filtered down to those where both genes are annotated as ‘gene with protein product’ in the HGNC (Braschi et al., 2018), and where the sequence identity is 20% in at least 1 direction of comparison. HGNC approved gene symbols and Entrez identifiers were also assigned to all paralogs using the HGNC ID mappings. We note that different releases of Ensembl define different paralog pairs, but found that Ensembl 93 paralog pairs had a greater overlap with the set of paralog pairs from the analysed combinatorial CRISPR screens in comparison to newer releases of Ensembl (e.g. Ensembl 102).

### Whole genome vs. small-scale duplicates

Paralog pairs are annotated as whole genome duplicates (WGDs) if they were included in the WGDs identified by (Makino and McLysaght, 2010) and/or the high-confidence (strict) list of WGDs in the OHNOLOGS v2 resource (Singh and Isambert, 2020). All other paralog pairs are classified as small-scale duplicates.

### Paralog pairs screened by Thompson et al

Thompson *et al*. screened a total of 1,191 gene pairs, including both paralog and non-paralog pairs, in three cell lines: A375 (melanoma), Mewo (melanoma) and RPE-1 (retinal epithelial) (Supplementary Data 1, (Thompson et al., 2021)). Of these gene pairs, 592 are paralog pairs that overlap with our reference list from Ensembl. To identify SL gene pairs (which we obtained from Thompson *et al*. Supplementary Data 5), Thompson *et al*. filtered out candidate pairs that contained a gene that had been identified as essential in the cell line(s) screened. We applied a similar filtering step for the non-hits; specifically, we filtered out all pairs that contain a gene that was identified as essential in all three cell lines screened (i.e. the gene did not pass any of the Mewo, A375 or RPE filters, see Thompson *et al*. Supplementary Data 1). This left us with a dataset of 475 paralog pairs for further analysis.

### Paralog pairs screened by Dede et al

Dede *et al*. screened 400 paralog pairs in three cell lines: A549 (lung cancer), HT29 (colorectal cancer) and OVCAR8 (ovarian cancer). Of these pairs, 393 overlap with our reference list from Ensembl. We obtained their table of zdLFC scores (Additional file 3:Table S2 (Dede et al., 2020)) and identified a pair as SL in a given cell line if it had zdLFC<-3, which is the threshold used by the authors. There are 24 pairs that fulfill this criterion in at least one of the screened cell lines.

### Paralog pairs screened by Parrish et al

Parrish *et al*. screened 1,030 paralog pairs in two cell lines: PC9 (lung adenocarcinoma) and HeLa (cervical carcinoma). Of these pairs, 1,027 overlap with our reference list from Ensembl. We obtained their genetic interaction (GI) scores (Supplemental Table 3 (Parrish et al., 2020)), and, as the authors did, considered a given paralog pair to be SL in a given cell line if it had GI score<-0.5 and GI FDR<0.1. There are 122 paralog pairs that fulfill these criteria in at least one of the two screened cell lines.

### Paralog pairs screened by Gonatopoulos-Pournatzis et al. (CHyMErA)

Gonatopoulos-Pournatzis *et al*. screened 688 paralog pairs in two cell lines: HAP1 and RPE-1. Of these pairs, 658 pairs overlap with our reference list of paralog pairs. We obtained their screen results (Table S8 (Gonatopoulos-Pournatzis et al., 2020)) and initially identified all pairs with a negative genetic interaction (GI) as reported by the authors; these are pairs whose ‘GI type’ is either ‘shared’, ‘unique_early’, or ‘unique_late’ and which have a negative effect size (i.e. pairs with a negative GI observed at any experimental time point). There are 267 paralog pairs that fulfill these criteria. We noted that not all of these negative GIs corresponded to synthetic lethality; in some cases the observed log-fold change (LFC) from the dual-gene knockout was close to zero, indicating that the combined gene loss was not detrimental to the cells. To identify SL pairs we filtered the negative GIs to those that have observed LFC<-0.9; we tested a range of LFC thresholds and found that filtering with -0.9 as the threshold resulted in the most significant overlap (based on Fisher’s exact test) between the CHyMErA SLs and the SLs from the three other combinatorial screens (Fig. S1A). With the addition of this LFC threshold we identify 72 (out of 267) paralog pairs as SL in at least one of the two screened cell lines. Considering all negative genetic interactions reported, the CHyMErA screen has a much higher hit rate (∼40.6%) than the three other screens, but after filtering for synthetic lethality its hit rate (∼11%) is consistent with that of the three other screens (Fig S1B).

### Filtering paralog pairs in our computational pipeline

To derive our dataset of SL paralog pairs, the reference set of paralog pairs obtained from Ensembl (see above) was filtered according to several criteria (Fig. 2A). We required paralog pairs to have at least 20% sequence identity in both directions of comparison (rather than just one) and filtered out pairs belonging to paralog families with more than 20 genes (to prevent over-representation of certain families). Family size was calculated as the length of the union of both genes’ (A1 and A2) sets of paralogs. We further filtered out paralog pairs where both genes are in general not “broadly expressed”. To determine this we obtained tissue distribution and specificity for mRNA expression from the Human Protein Atlas (version 19.3, file: proteinatlas.tsv) (Uhlén et al., 2015). We consider genes to be “broadly expressed” if their expression is detected in all tissues, or if their expression is detected in at least a third of tissues (‘detected in many’) and their tissue specificity is classified as low or not detected.

### Gene dependency scores from DepMap (Broad)

We reprocessed the DepMap screen data for 769 cancer cell lines (DepMap Release 20Q2, file:Achilles_logfold_change.csv) with CERES, which corrects for the known effects of copy number alterations (Meyers et al., 2017), after dropping multi-targeting single-guide RNAs (Table S2). Multi-targeting guides were identified by the same methodology as in (De Kegel and Ryan, 2019) and include guides that align to multiple protein-coding genes with either a perfect match, a single mismatch, or a PAM-distal double mismatch. Only scores for protein-coding genes that were targeted by at least three unique guides are retained for further analysis. Overall 1,681 of the 18,119 genes with CERES scores in DepMap were dropped from our analysis (some of these genes were dropped simply because they are not classified as protein-coding by the HGNC).

### Calling gene loss from expression, mutation and copy number data

We obtained RNA-seq (file:CCLE_expression.csv), gene mutation (file:CCLE_mutations.csv) and gene level copy number (file:CCLE_gene_cn.csv) data from the DepMap portal (release 20Q2) (Ghandi et al., 2019). Data from all three sources was available for 762 of the 769 cell lines included in the DepMap CRISPR screens, so gene loss was identified for these 762 cell lines (Table S3). A gene is marked as lost in a given cell line if at least one of the following conditions is met: gene expression (log TPM+1) < 1, gene expression z-score < -4, the gene has a LOF mutation (splice site, nonsense, or frameshift), or the gene is homozygously deleted. We consider a gene to be homozygously deleted if its copy number ratio is < -1.28 and the gene’s homozygous deletions are in general supported by a relative decrease in expression. Specifically, we test that the gene’s expression is significantly lower in the cell lines where it is homozygously deleted, compared to the cell lines where it is not (t-test *p*<0.05).

### Identifying putative synthetic lethal paralog pairs

To identify a significant association between A2 gene loss and A1 dependency we used a model with the form: A1_dependency ∼ A2_status + C(cell_line_lineage). Cell line lineage information was obtained from the DepMap portal (release 20Q2, file: sample_info.csv). Pairs where the A1 gene was not essential (essential = CERES score < -0.6) in any of 762 cell lines for which we had data were directly marked as non-SL. For the remaining pairs we applied Benjamini- Hochberg multiple testing correction (Benjamini and Hochberg, 1995) on the *p*-value for the A2_status variable. Pairs with a negative A2_status coefficient were considered SL at a false discovery rate (FDR) of 10%.

### Gene dependency scores from Behan et al

We obtained the log-fold change data from the Behan *et al*. genome-wide CRISPR screens of 318 cell lines (Behan et al., 2019) from the DepMap portal (Sanger CRISPR (CERES), file:logfold_change.csv). As for the Broad DepMap screen data (see above), we processed these log-fold changes with CERES after dropping multi-targeting single-guide RNAs. Of the 318 screened cell lines, 242 overlap with the CCLE molecular profiling data and these are used in the validation of our dataset of SL and non-SL pairs (Fig. 2C).

### Budding and fission yeast orthologs

*S. pombe* orthologs were obtained from PomBase (Lock et al., 2019) and their essentiality status (essential vs. non-essential) was obtained from (Kim et al., 2010) (Supplementary Table S1). *S. cerevisiae* orthologs were downloaded from Ensembl v93 (Zerbino et al., 2018) and their essentiality status was obtained from OGEE v2 (Chen et al., 2017). Orthologs without available essentiality data were marked as non-essential because the majority of yeast genes are non-essential (Gu et al., 2003; Kim et al., 2010).

### Conservation score

The conservation score of a paralog pair was calculated as the number of species, out of nine, in which at least one member of the pair has an identifiable ortholog. The nine species considered are: *Arabidopsis thaliana, Caenorhabditis elegans, Danio rerio, Drosophila melanogaster, Escherichia coli, Mus musculus, Rattus norvegicus, S. cerevisiae* and *Xenopus tropicalis*. Orthologs were obtained from Ensembl v93 (Zerbino et al., 2018) with the exception of *A. thaliana* and *E. coli* orthologs which were not available in Ensembl v93 and were instead obtained from InParanoid (Sonnhammer and Östlund, 2015).

### Gene age

Gene age for each individual gene in the paralog pairs was obtained from ProteinHistorian (Capra et al., 2012), with the default age estimation options (Family database = PPODv4_PTHR7-OrthoMCL, Reconstruction algorithm = wagner1.0). Although the genes in a paralog pair are theoretically expected to have the same age, discrepancies in the gene trees used to determine age and paralogy can lead to different age estimates. Consequently we used the average age of the two paralogs as the age for the pair.

### Protein-protein interactions

Protein-protein interactions were obtained from BioGRID (version 3.5.187) (Oughtred et al., 2019). The full BioGRID data set was filtered for physical interactions between two human protein-coding genes. Genes in a paralog pair are annotated as interacting if there is any experimental evidence of an interaction between them. In general we use all non-redundant interactions, i.e. we disregard directionality, experimental system or publication when counting interactions.

### Protein complex membership

Sub-units for known protein complexes were obtained from CORUM (Giurgiu et al., 2019) (complete complexes dataset). A paralog pair was labelled as a protein complex member if at least one of its member genes is annotated as coding for a protein complex sub-unit.

### Gene expression features

Gene expression in 17,382 non-diseased tissue samples was obtained from the GTeX database (GTEx Consortium, 2020). For each gene in each paralog pair we computed the mean expression over all samples. This was used to populate two features for each pair: mean expression (lower), i.e. the mean of the lower expressed gene in the pair, and mean expression (higher), i.e. the mean of the higher expressed gene in the pair. We also computed Spearman’s correlation between the expression of the two genes in each paralog pair across all samples.

### Subcellular location

Subcellular location data was obtained from the Human Protein Atlas (version 19.3, file: subcellular_location.tsv, (Uhlén et al., 2015)). We used both the main and additional locations associated with each gene, excluding annotations with “Uncertain” reliability.

### Feature value imputation

For eight of our 22 features data was not available for all paralog genes and/or pairs. As the random forest model we used does not allow missing feature values, these features were set to either zero or the mean value across all paralog pairs. For shared domains and co-localisation data wasn’t available for ∼2.94% and ∼41.76% of all paralog genes respectively; as these features are the Jaccard index their values are zero in case data wasn’t available for both or either of the genes. Imputation affects only a small percentage of paralog pairs; specifically, the features and associated percentages of pairs affected are: shared PPI mean essentiality (∼1.3% of pairs with shared PPIs), mean complex essentiality (∼0.74% of pairs in protein complexes), mean expression (higher and lower) (∼0.23% of all pairs), expression correlation (∼1.38% of all pairs) and gene age (∼4.13% of all pairs). The mean complex essentiality for pairs that don’t include a protein complex member, and the mean essentiality of shared PPIs for pairs that do not have any shared PPIs were filled in with zero.

### Metrics for individual features

For each feature we computed the ROC AUC and average precision for the feature values as well as the inverse of the feature values (i.e. we considered both a positive and negative relationship between the feature and synthetic lethality). We then retained the maximum ROC AUC and average precision values. For all features except family size and expression correlation the maximum metric values were observed for a positive relationship between the feature and synthetic lethality.

### The random forest classifier

We use the random forest classifier implementation in the scikit-learn package (class RandomForestClassifier). We performed a grid search based on stratified cross-validation of our paralog pair dataset to explore the space of potential hyper-parameters for the random forest classifier. To prevent overfitting we select relatively low maximum tree depth and relatively high minimum leaf node weight. The parameters used in the final model are as follows: n_estimators=600 (i.e. number of trees), max_features=0.5, max_tree_depth=3, min_child_weight=8, and random_state=8 (for reproducibility). All parameters not listed are left as the default specified in scikit-learn (version 0.23.1).

### SL predictions from SLant

We obtained SLant predictions for an extensive list of gene pairs via personal communication with the authors (Benstead-Hume et al., 2019). This is an extended version of what is available in the Slorth database, which only includes scores for those pairs that were predicted to be SL (i.e. pairs with a prediction score above a certain threshold).

### SL predictions from SINaTRA

We obtained the full set of SINaTRA prediction scores for human gene pairs from three files (<u>10.6084/m9.figshare.1501103</u>, <u>10.6084/m9.figshare.1501105</u>, <u>10.6084/m9.figshare.1501115</u>) provided with the paper (Jacunski et al., 2015).

### SL predictions from Cai et al

We generated the predictions from the classifier developed by Cai *et al*. by running the case_study.py script from the associated github repository (https://github.com/CXX1113/Dual-DropoutGCN) with parameter TOP_K set to ’all’ and all other parameters left as the default. This script trains their classifier (a dual-dropout graph convolutional network) using their full training dataset and predicts novel SLs among all the unknown gene pairs.

### Feature value z-scores

For numeric features we compute z-scores using feature values for all ∼36.6k paralog pairs with at least 20% sequence identity in at least one direction of gene-to-gene comparison (Ensemble reference set). These z-scores were then capped at +/-2. For Boolean features the ‘z-scores’ are simply set to 2 and -2 for True and False feature values respectively, i.e. we don’t take the ratio of True/False into account.

### siRNA experiments

HAP1 cell lines were obtained from Horizon Discovery: HAP1_WT (C631), HAP1_COPS7A_ KO (HZGHC000914c001), HAP1_COPS7B_KO (HZGHC000636c011), HAP1_ASF1A_KO (HZGHC007215c005) and HAP1_ASF1B_KO (HZGHC55723). Cells were cultured in IMDM (10-016-CVR; Corning) supplemented with 10% FBS (10270-106; Thermo Fisher Scientific). ON-TARGETplus siRNA SMARTpools targeting COPS7A (L-020873-01-0005), COPS7B (L- 014207-02-0005), PLK1 (L-003290-00-0005), ASF1A (LQ-020222-02-0002), ASF1B (LQ-020553-00-0002) and a non-targeting scramble control (D-001810-10-20) were obtained from Dharmacon. HAP1 cells were seeded to a density of 5000 cells per well of a 96-well plate and 5nM siRNA was transfected with Lipofectamine 3000 (L3000015; Thermo Fisher Scientific) in Opti-MEM I Reduced Serum Medium (31985070; Thermo Fisher Scientific). Cell viability was measured 72 hours after siRNA transfection using CellTiter-Glo Luminescent Cell Viability Assay (G7570; Promega). The 96 well plates were read using a SpectraMax® M3 Microplate Reader (Molecular devices). Viability effects for each siRNA targeting each gene X in each cell line y were normalised using the following formula:

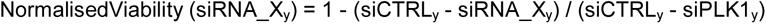

where siCTRL_y_ is the average of 3 measurements of non-targeting scramble control in cell line y and siPLK1_y_ is the average of 3 measurements of an siRNA smartpool targeting PLK1 in cell line y. Raw and normalised viability data are in Table S10.

## Acknowledgements

B.D.K., N.Q. and C.J.R. were funded through an Irish Research Council 2017/2018 Laureate Award awarded to C.J.R.. D.J.A. and N.A. were funded from the Wellcome Trust, Cancer Research UK and the European Research Council under the European Union’s Seventh Framework Programme (FP7/2007–2013) / ERC synergy grant agreement n° 319661 COMBATCANCER. We thank Graeme Benstead-Hume and Frances Pearl for sharing the complete list of predictions generated by SLant and Traver Hart for sharing screen data ahead of publication. We thank Barry Scott for the introduction to SHAP values and Ishan Mehta and Chris Lord for helpful feedback.

## Supplemental Figures

**Figure S1.**
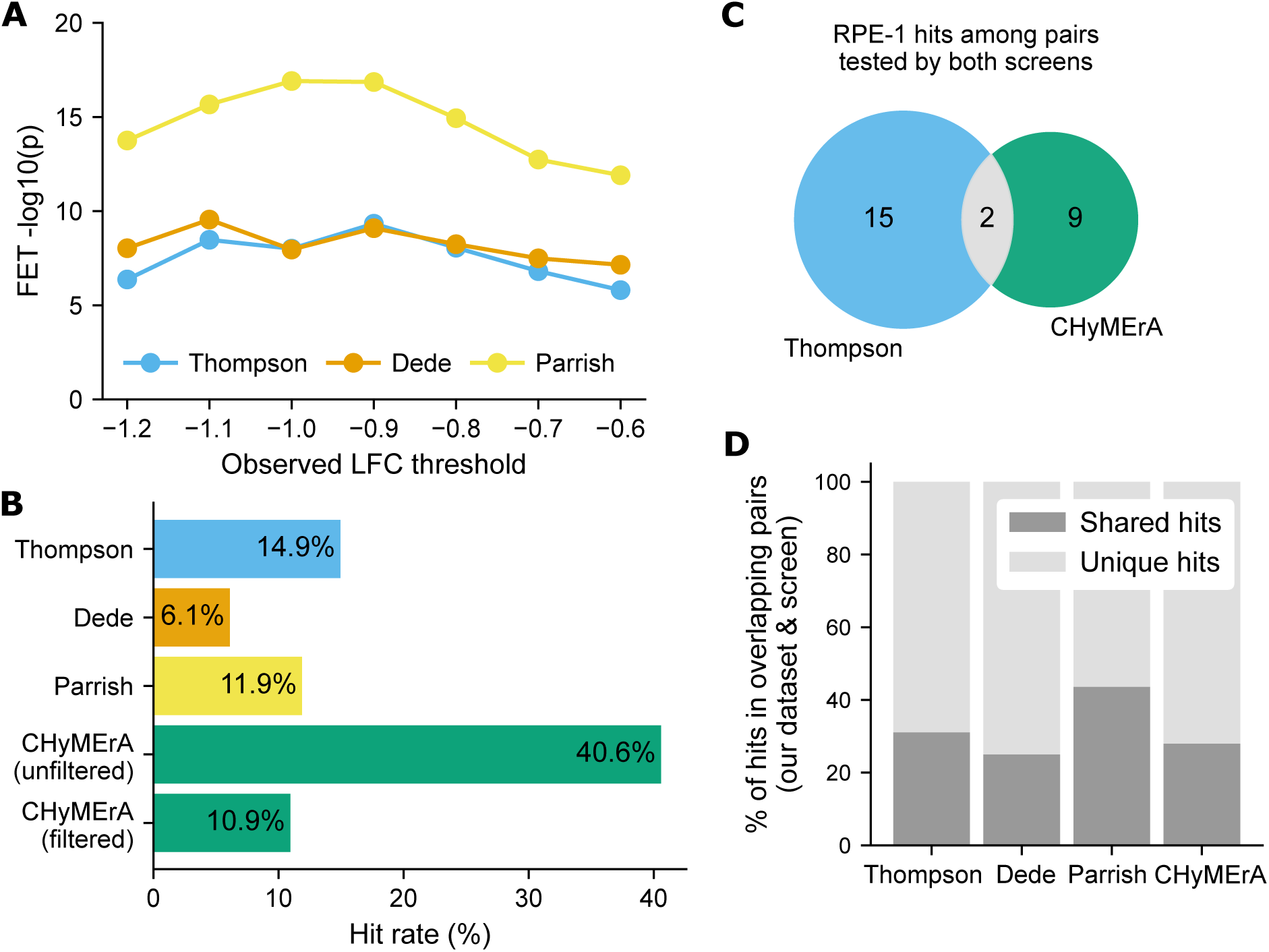
(A) For each of the Thompson, Dede and Parrish combinatorial screens, the FET -log10 *p*-value (y-axis) for the overlap of hits in that screen and hits in the CHyMeRA SL subset obtained from filtering with the indicated CHyMeRA double knockout LFC thresholds (x-axis). (B) Bar chart showing the percentage of tested paralog pairs that are identified as hits; for “CHyMeRA (unfiltered)” hits correspond to negative genetic interactions, in all other cases hits correspond to SL interactions. (C) The overlap of RPE-1 hits identified in the Thompson and CHyMeRA screens, among paralog pairs tested in both screens. (D) For the pairs that are both in each screen and our dataset, the percentage of hits that are shared vs. unique to either the screen or our dataset.

**Figure S2.**
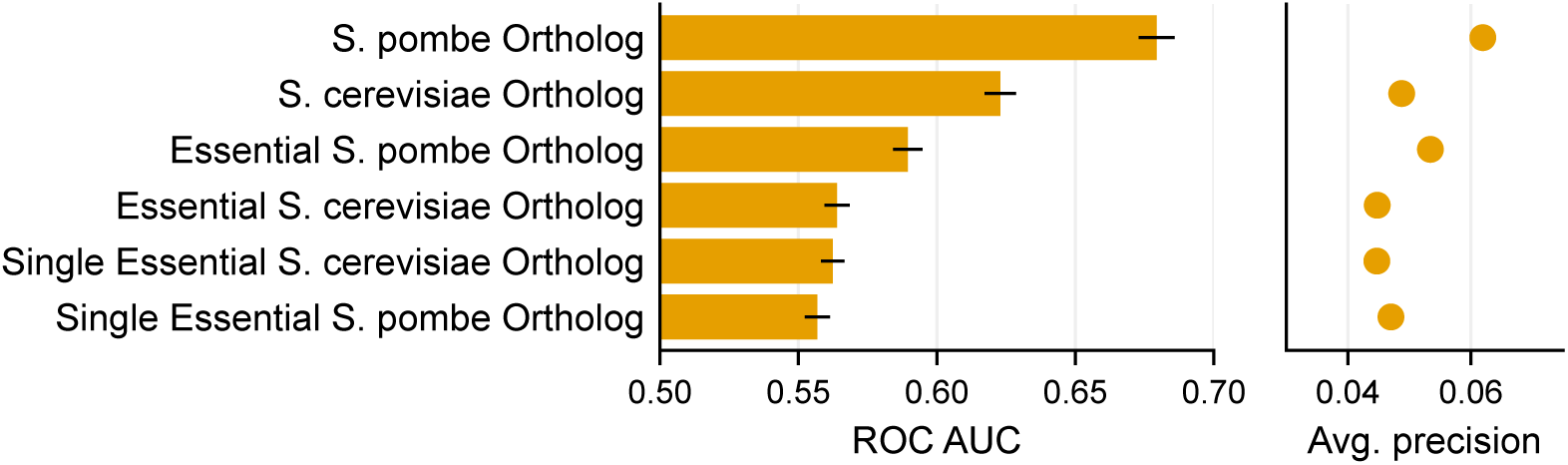
Bar chart showing the area under the ROC curve (ROC AUC) and point plot showing the average precision for six budding and fission yeast features treated as individual classifiers. The error bars show the mean standard error based on computing the ROC AUC and average precision for 10 randomly selected, stratified fourths of our dataset.

**Figure S3.**
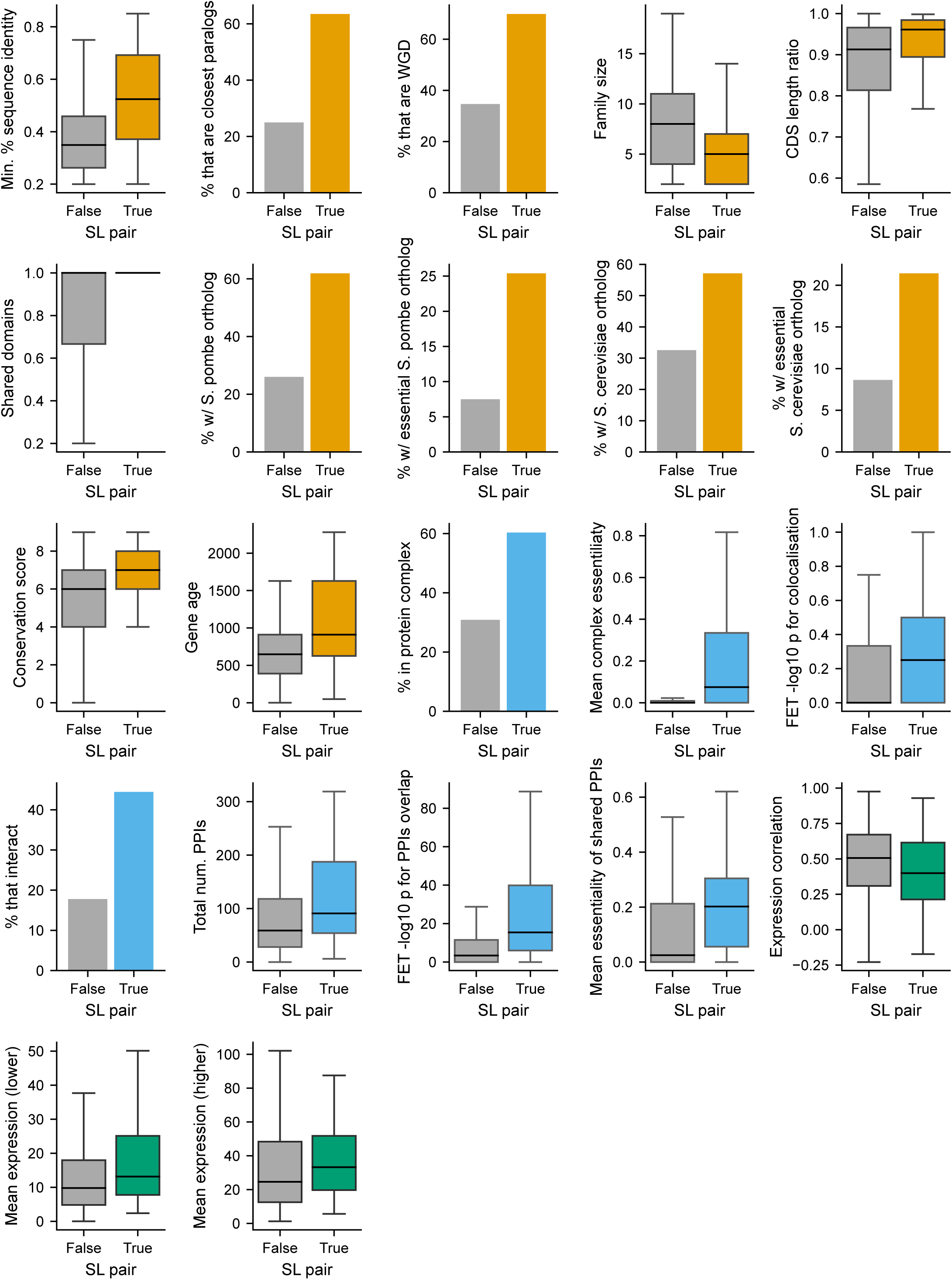
Boxplots for each quantitative feature and bar charts for each Boolean feature showing the difference in feature distributions for SL and non-SL paralog pairs in our dataset. The features are the same as those shown in Fig. 3A. For each box plot, the black central line represents the median, the top and bottom represent the 1st and 3rd quartile, and the whiskers extend 1.5 times the interquartile range past the box. Outliers are not shown.

**Figure S4.**
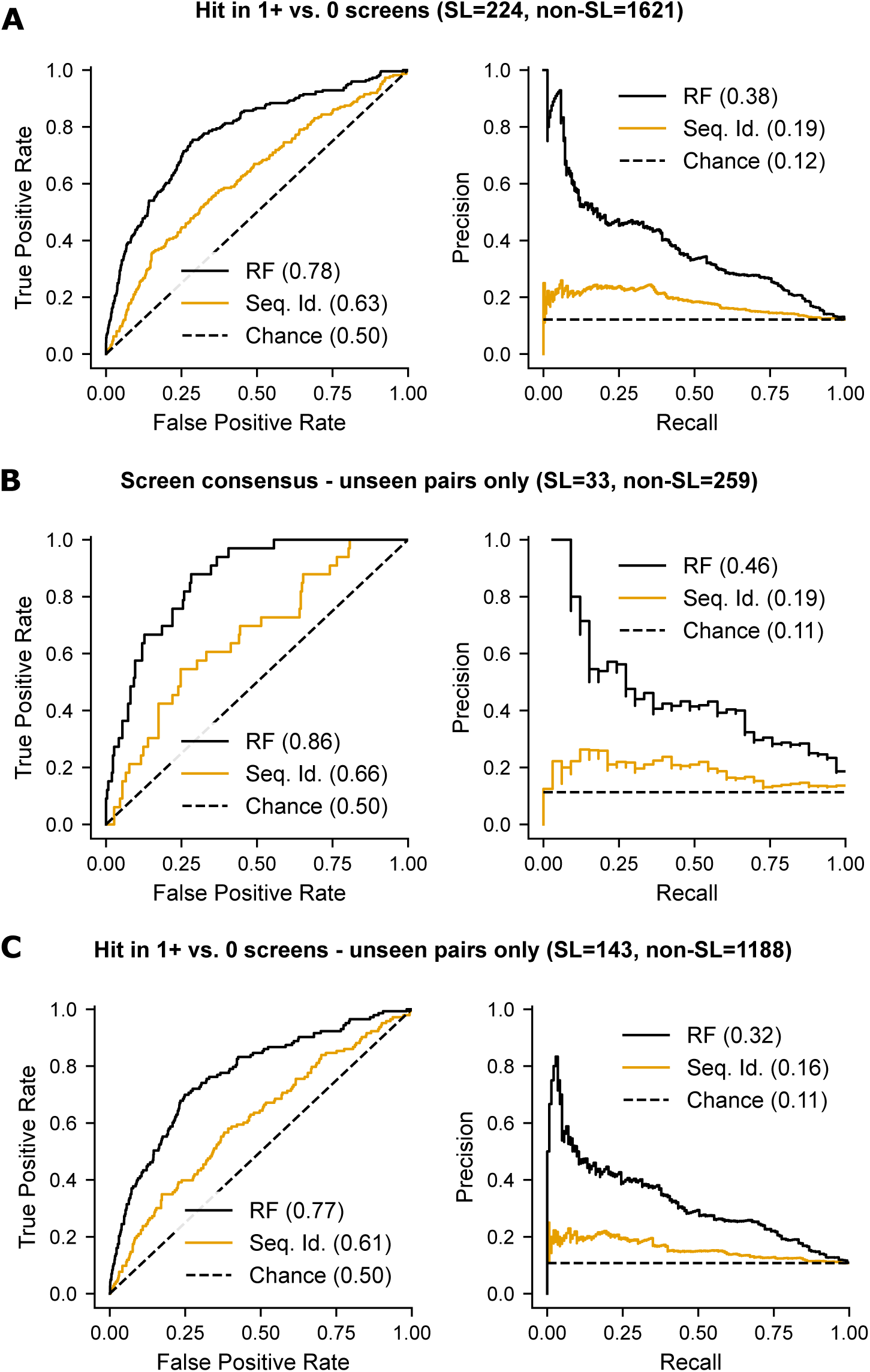
(A) ROC and PR curves showing the performance of our classifier (black) and sequence identity (orange) in distinguishing paralog pairs SL in 1+ vs. 0 combinatorial screens. The numbers shown in the legends are ROC AUC and average precision. (B) Same as Fig. 4A (distinguishing screen consensus SLs vs. non-SLs) but for just the subset of screened paralog pairs that are not in our training data. (C) Same as A but for just the subset of screened paralog pairs that are not in our training data.

**Figure S5.**
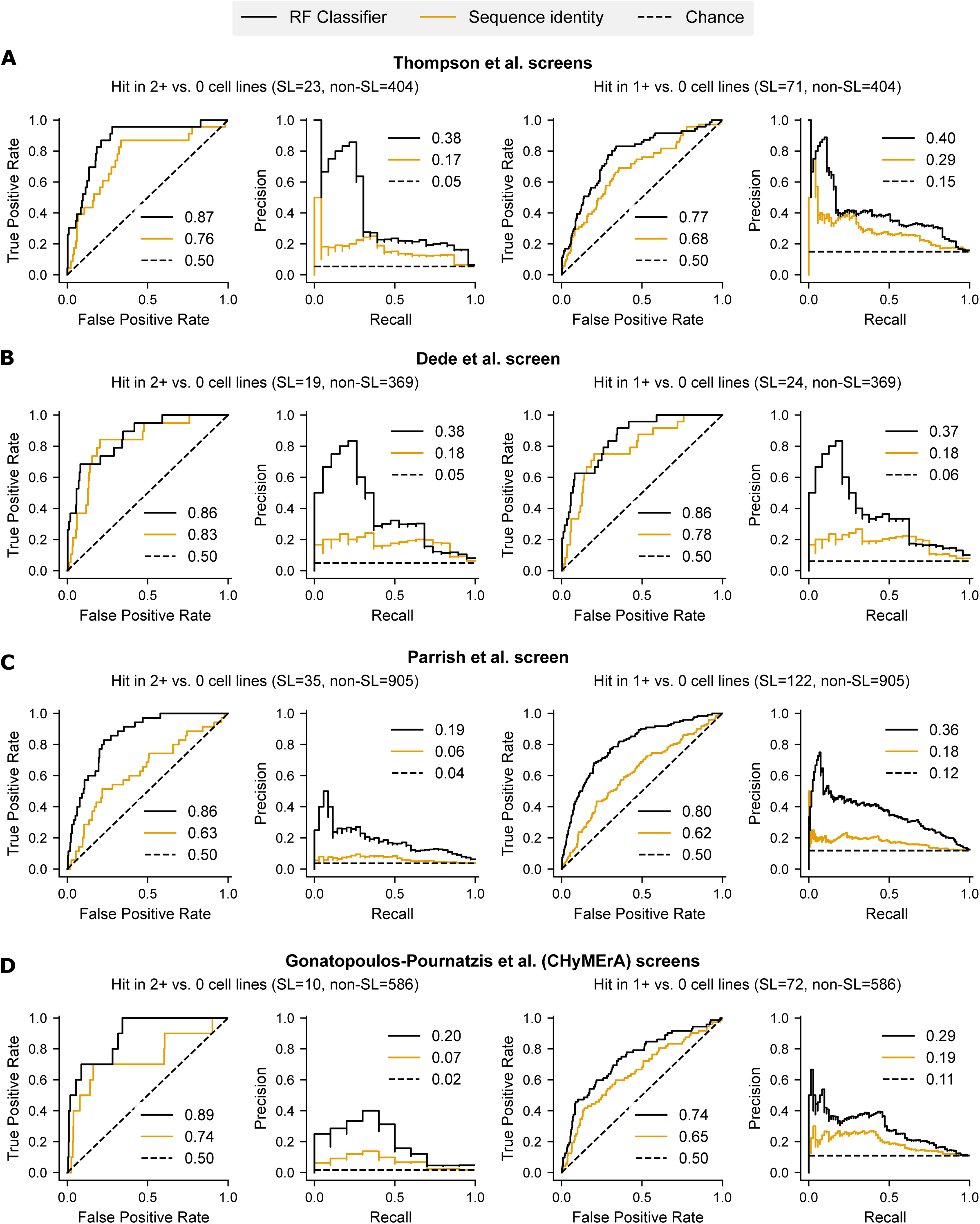
ROC and PR curves showing the performance of our classifier (black) and sequence identify (orange) in distinguishing paralog pairs that were found to be SL in 2+ vs. 0 cell lines (left) or 1+ vs. 0 cell lines (right) in each of four combinatorial screens: (A) Thompson *et al*., (B) Dede *et al*., (C) Parrish *et al*., and (D) Gonatopoulos-Pournatzis *et al*. (CHyMErA). The legends for the ROC plots show the ROC AUC for each curve while the legends for the PR plots show the average precision associated with each curve.

**Figure S6.**
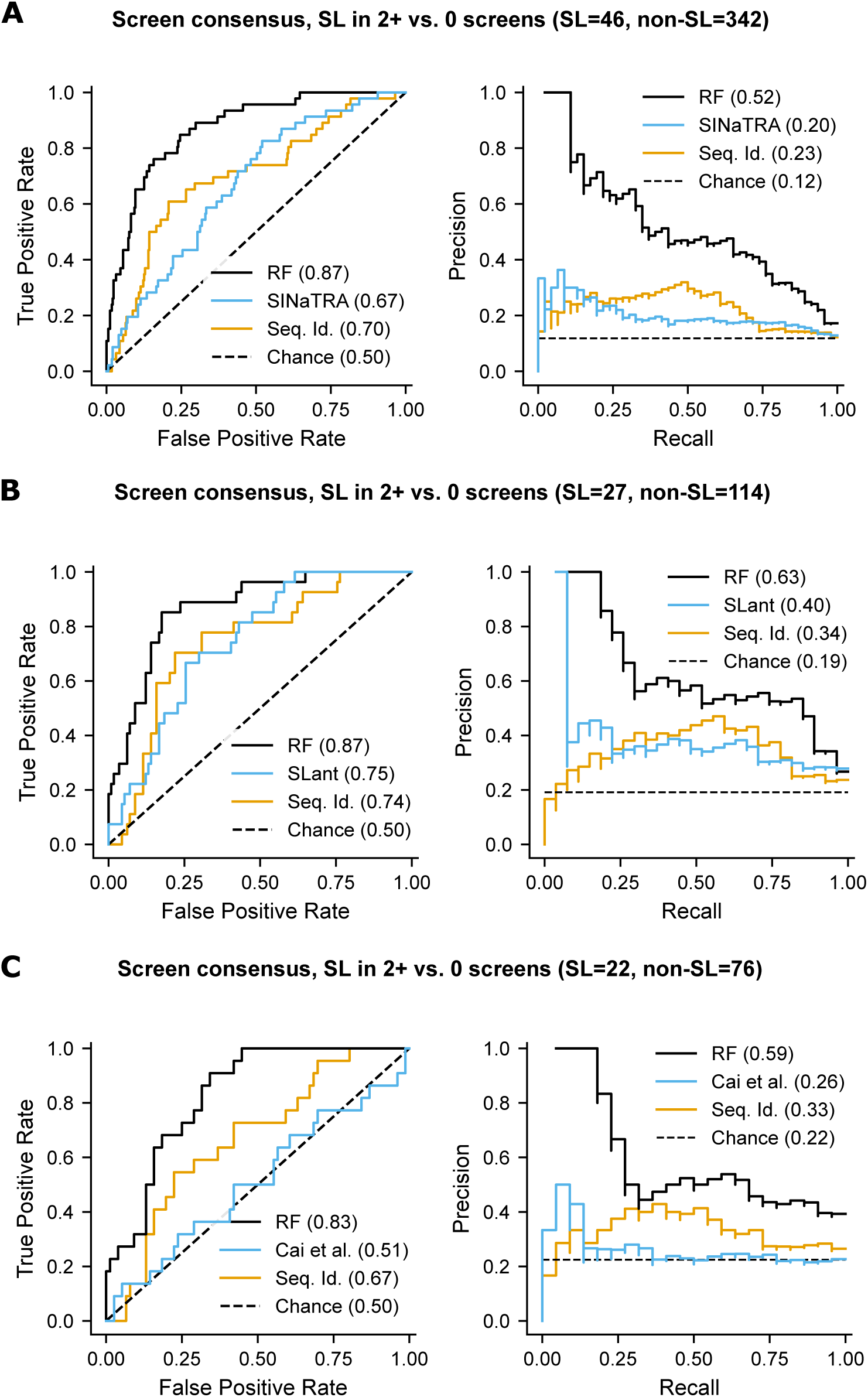
(A) ROC and PR curves showing the performance of the SINaTRA SL classifier (blue), our random forest classifier (black) and sequence identity (orange) in distinguishing paralog pairs that were identified as SL in 2+ vs. 0 (and screened in 2+) combinatorial screens. The performance comparison was made for the subset of screened paralog pairs for which SLant predictions were available (numbers shown in the graph title). The numbers shown in the legends are ROC AUC and average precision. (B) Same as (A) but for the SLant classifier. (C) Same as (A) but for the Cai *et al*. classifier.

**Figure S7.**
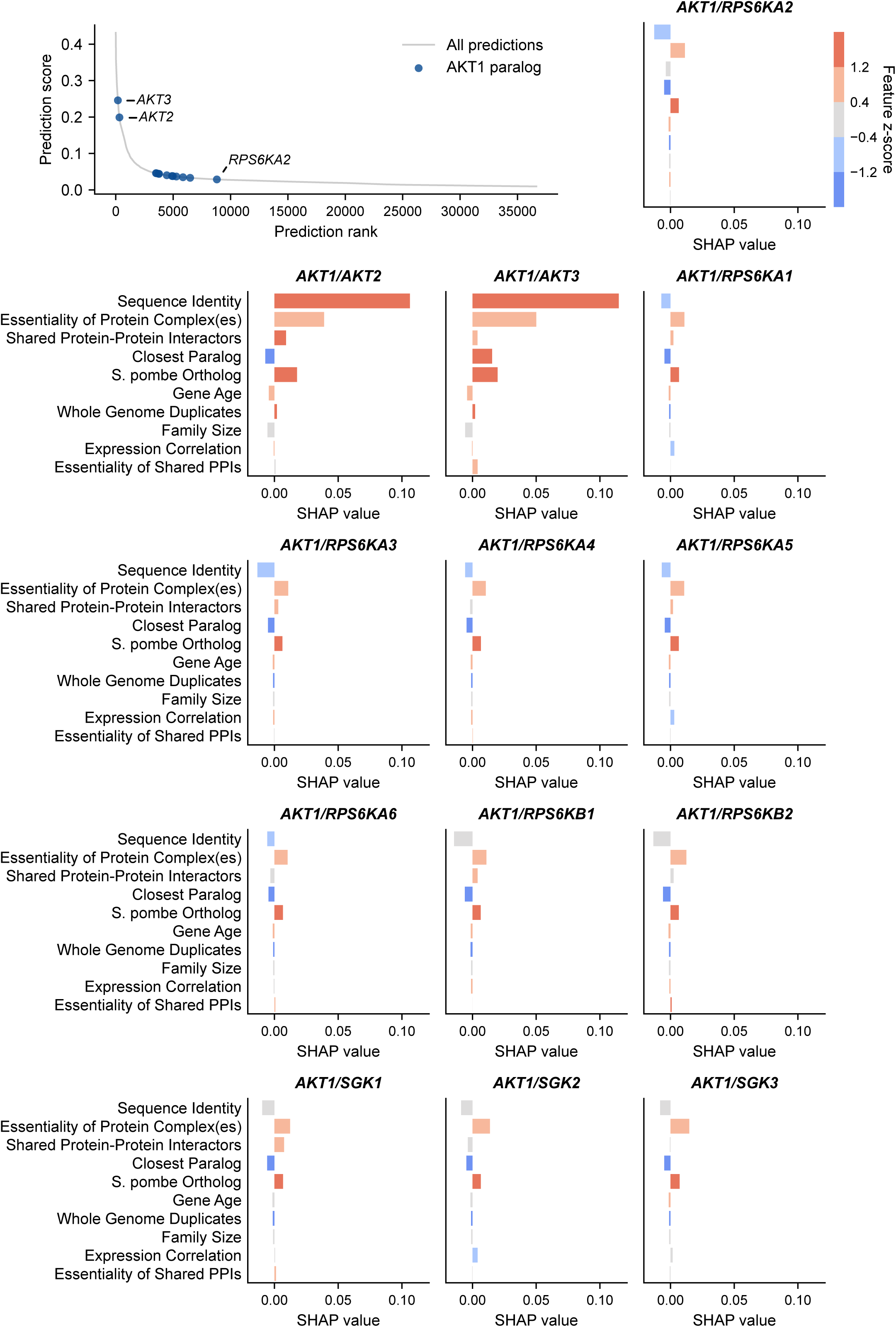
Top left: Plot showing the prediction score vs. rank for all paralog pairs (as in Fig. 5A) with paralog pairs that contain the gene *AKT1* highlighted in dark blue. Top right/bottom: SHAP profiles for all paralog pairs that contain *AKT1*.

## Supplemental Files

**Table S1.** Paralog synthetic lethalities with low throughput experimental validation curated from the literature.

**Table S2.** Gene dependency scores computed by CERES across 769 cell lines, after filtering out potential multi-targeting guides. Columns are gene Entrez IDs, rows are DepMap cell line IDs.

**Table S3.** Binary matrix that indicates, for each gene (A2 in the paralog pairs in Table S4) in each cell line, whether the gene is lost (1) or not (0) in that cell line. Rows are Entrez gene IDs and columns are DepMap cell lines IDs.

**Table S4.** Results of testing 3,810 candidate paralog pairs for an association between A1 dependency and A2 loss. Pairs are listed twice if they were tested symmetrically, i.e. their genes were considered as the A1 and A2 gene in turn. The A2_status_coef, A2_status_p and A2_status_p_adj columns provide, respectively, the coefficient of the A2_status variable, the raw p-value and the FDR corrected p-value for the association between A1 dependency and A2 loss for each pair. Pairs for which the SL field is blank (NaN) were dropped from further analysis.

**Table S5.** Final dataset of SL and non-SL paralog pairs annotated with 22 paralog pair features. These pairs are a subset of those in Table S4.

**Table S6.** Descriptions of all 22 paralog pair features and the ROC AUC and average precision obtained from treating each feature as an individual classifier for our dataset (Table S5). For quantitative features: the *p*-value from a two-sided Mann-Whitney test, and for Boolean features: the *p*-value from a Fisher’s exact test (FET) comparing SL and non-SL pairs from our dataset.

**Table S7.** ROC AUC and avg. precision (AP) values for four test datasets for three previously developed classifiers (Other = SINaTRA, SLant and Cai et al. classifiers) compared to our classifier (RF) and sequence identity (Seq_id). The test datasets are: the screen consensus dataset (hits in 2+ vs. 0 screens), all screen hits (hits in 1+ vs. 0 screens) and the versions of those with unseen paralog pairs only (i.e. pairs not included in our training data). The number of SL and non-SLs in each dataset for which predictions from the other classifier were available are shown. ROC AUC and AP values are rounded to two decimal places.

**Table S8.** Feature values and predictions (rank, percentile and score) for all ∼36.6k paralog pairs. Table also indicates whether the pair has been validated as SL (i.e. is in Table S1), the number of combinatorial screens in which the pair was tested (n_screens) and identified as SL (n_screens_SL) as well as whether the pair was a hit in our computational dataset (depmap_hit).

**Table S9.** SHAP values associated with all features for all predictions from Table S8.

**Table S10.** ASF1A/ASF1B and *COPS7A/COPS7B* viability data.

